# Potent and broadly neutralizing antibodies against sarbecoviruses elicited by single ancestral SARS-CoV-2 infection

**DOI:** 10.1101/2024.06.06.597720

**Authors:** Lei Yu, Yajie Wang, Yuanchen Liu, Xiaomin Xing, Chen Li, Xun Wang, Jialu Shi, Wentai Ma, Jiayan Li, Yanjia Chen, Rui Qiao, Xiaoyu Zhao, Ming Gao, Shuhua Wen, Yingxue Xue, Yongjun Guan, Hin Chu, Lei Sun, Pengfei Wang

**Affiliations:** Guangzhou Eighth People’s Hospital, Guangzhou Medical University, Guangzhou, China; Shanghai Fifth People’s Hospital, Shanghai Institute of Infectious Disease and Biosecurity, Institutes of Biomedical Sciences, Fudan University, Shanghai, China; State Key Laboratory of Emerging Infectious Diseases, Department of Microbiology, School of Clinical Medicine, Li Ka Shing Faculty of Medicine, The University of Hong Kong, Pokfulam, Hong Kong Special Administrative Region, China; Shanghai Pudong Hospital, Fudan University Pudong Medical Center, State Key Laboratory of Genetic Engineering, MOE Engineering Research Center of Gene Technology, School of Life Sciences, Shanghai Institute of Infectious Disease and Biosecurity, Fudan University, Shanghai, China; Beijing Institute of Genomics, Chinese Academy of Sciences, University of Chinese Academy of Sciences and China National Center for Bioinformation, Beijing, China; Antibody BioPharm, Inc., Gaithersburg, MD, USA; Changyuan Funeng (Shanghai) Life Technology Co., Ltd., Shanghai, China

## Abstract

Monoclonal antibody (mAb) therapeutics hold promise for both preventing and treating infectious diseases, especially among vulnerable populations. However, the emergence of various variants of severe acute respiratory syndrome coronavirus 2 (SARS-CoV-2) presents challenges for current mAb treatments, emphasizing the need for more potent and broadly neutralizing antibodies. In this study, we employed an unbiased screening approach to discover broadly neutralizing antibodies and successfully isolated two mAbs from individuals with only exposure to ancestral SARS-CoV-2. One of these antibodies, CYFN1006-1, exhibited robust cross-neutralization against a spectrum of SARS-CoV-2 variants, including the latest JN.1 and KP.2 variants, with consistent IC_50_ values ranging from ∼1 to 5 ng/mL. Notably, it also displayed broad neutralization activity against SARS-CoV and related sarbecoviruses, such as WIV1, SHC014, RaTG13, and GD-Pangolin. Structural analysis revealed that these mAbs target shared hotspot but mutation-resistant epitopes, with their Fabs locking the RBD in the “down” conformation through interactions with adjacent Fabs and RBDs, and cross-linking Spike trimers into di-trimers to block viral infection. In vivo studies conducted in a JN.1-infected hamster model validated the protective efficacy of CYFN1006-1, emphasizing its therapeutic potential. These findings suggest that, through meticulous approaches, rare antibodies with cross-neutralization activities against SARS-CoV-2 and related sarbecoviruses can be identified from individuals with exclusively ancestral virus exposure.

## Main

The relentless spread of severe acute respiratory syndrome coronavirus 2 (SARS-CoV-2) has led to millions of infections and substantial morbidity and mortality across continents, severely challenging healthcare systems, economies, and societal norms. The clinical spectrum of SARS-CoV-2 infection ranges from asymptomatic or mild respiratory illness to severe pneumonia, acute respiratory distress syndrome, multi-organ failure, and death, with certain populations, such as the elderly and those with underlying health conditions, being at higher risk of severe outcomes^1^. The genome of SARS-CoV-2 encodes several structural and non-structural proteins crucial for viral replication, transcription, and immune evasion. Among these proteins, the Spike (S) protein plays a pivotal role in viral entry by binding to the angiotensin-converting enzyme 2 (ACE2) receptor on host cells, facilitating membrane fusion, and initiating infection^2, 3^. The S protein is a major target for host immune responses and serves as the primary antigen for vaccine development and immunotherapeutic interventions.

Monoclonal antibody (mAb) therapeutics targeting the S protein of SARS-CoV-2 represent a vital option for both prophylaxis and treatment, especially for individuals who are immunocompromised, unable to receive vaccination, or have other risk factors predisposing them to severe disease^4^. Throughout the course of infection, antibodies exert their therapeutic effects by various mechanisms, including neutralizing virions through blocking receptor binding, crosslinking viral proteins, inhibiting fusion with host cells, and facilitating immune clearance of virions and infected cells^5^.

However, as SARS-CoV-2 continues to evolve, numerous variants carrying multiple mutations have emerged, resulting in significant evasion of antibody neutralization^6–9^. All anti-SARS-CoV-2 mAbs and mAb cocktail therapeutics previously approved for Emergency Use Authorization (EUA) by the FDA were subsequently withdrawn due to immune escape^10^, except for one exception—PEMGARDA (pemigard), which recently obtained EUA for pre-exposure prophylaxis (PrEP)^11^. Some early-discovered antibodies and their combinations, such as Bamlanivimab/Etesevimab, Casirivimab/imdevimab (REGN-COV2), and Tixagevimab/Cilgavimab (Evusheld), lost their neutralizing activity against SARS-CoV-2 variants^9, 12–16^. Bebtelovimab (LY-CoV1404), while highly efficient in neutralizing many SARS-CoV-2 variants, including early Omicron subvariants, escaped neutralization by subsequently emerging BQ.1.1 and XBB lineages^15, 17^. Another mAb isolated from a SARS-CoV convalescent individual, Sotrovimab (S309), maintained its neutralizing activity against most emerged SARS-CoV-2 variants due to recognition of a conserved epitope^4, 8, 18, 19^. However, its neutralizing potency is mediocre, with an IC_50_ value around 1 μg/ml against many SARS-CoV-2 variants^15, 19^, leading to its EUA withdrawal as well. These findings underscore the imperative for the development of next-generation antibodies capable of effectively resisting immune evasion induced by SARS-CoV-2 evolution.

To date, some neutralizing antibodies with cross-neutralization activities against SARS-CoV-2 and its variants have been identified. One example is SA55, isolated from SARS-CoV-2-vaccinated SARS convalescents has been reported to effectively resist mutations that have emerged in SARS-CoV-2^20^. Additionally, antibodies targeting the SD1 region, such as S3H3 and sd1.040, have demonstrated efficient cross-neutralization against major SARS-CoV-2 variants due to sequence conservation^21–23^. While the S2 subunit exhibits greater conservation compared to S1, antibodies targeting S2 typically exhibit suboptimal potency for achieving protection^4, 24–26^. Recently, we reported the isolation of potent and broadly neutralizing antibodies from a vaccinated donor who received a special five-dose COVID-19 vaccination schedule. Among these, two mAbs, PW5-5 and PW5-535, were found to possess pan-sarbecovirus potential^27^. However, since most of these broadly neutralizing antibodies were isolated from individuals who received multiple vaccinations or experienced multiple infections, or a combination of both, it remains unclear whether a single infection with the prototype strain could elicit antibodies capable of broadly neutralizing mutant viruses to which the donor had not been previously exposed.

### SARS-CoV-2 and SARS-CoV cross-reactive antibodies from ancestral SARS-CoV-2 infection

We recruited three COVID-19 convalescents who had been infected with the original SARS-CoV-2 virus during the initial outbreak in China, a period when vaccines were not yet available (**Fig. 1a**). All participants had confirmed viral exposure, as evidenced by positive RNA tests, and had received treatment at Guangzhou Eighth People’s Hospital. In May 2020, four months after their infection, we collected peripheral blood mononuclear cells (PBMCs) from these individuals. Memory B cells (MBCs) were subsequently sorted, and all MBCs were cultured without bias towards any specific antigen (**Extended Data Fig. 1**). The responses of SARS-CoV-2 and SARS-CoV cross-reactive MBCs have been investigated, and the findings are forthcoming in another publication. From a single well of these cultured MBCs from Pt2, which demonstrated cross-reactivity to both SARS-CoV-2 and SARS-CoV, we cloned one antibody heavy chain and two light chains (**Fig. 1b**), which were then assembled into two human IgG1 antibodies, named CYFN1006-1 and CYFN1006-2.

**Fig. 1:**
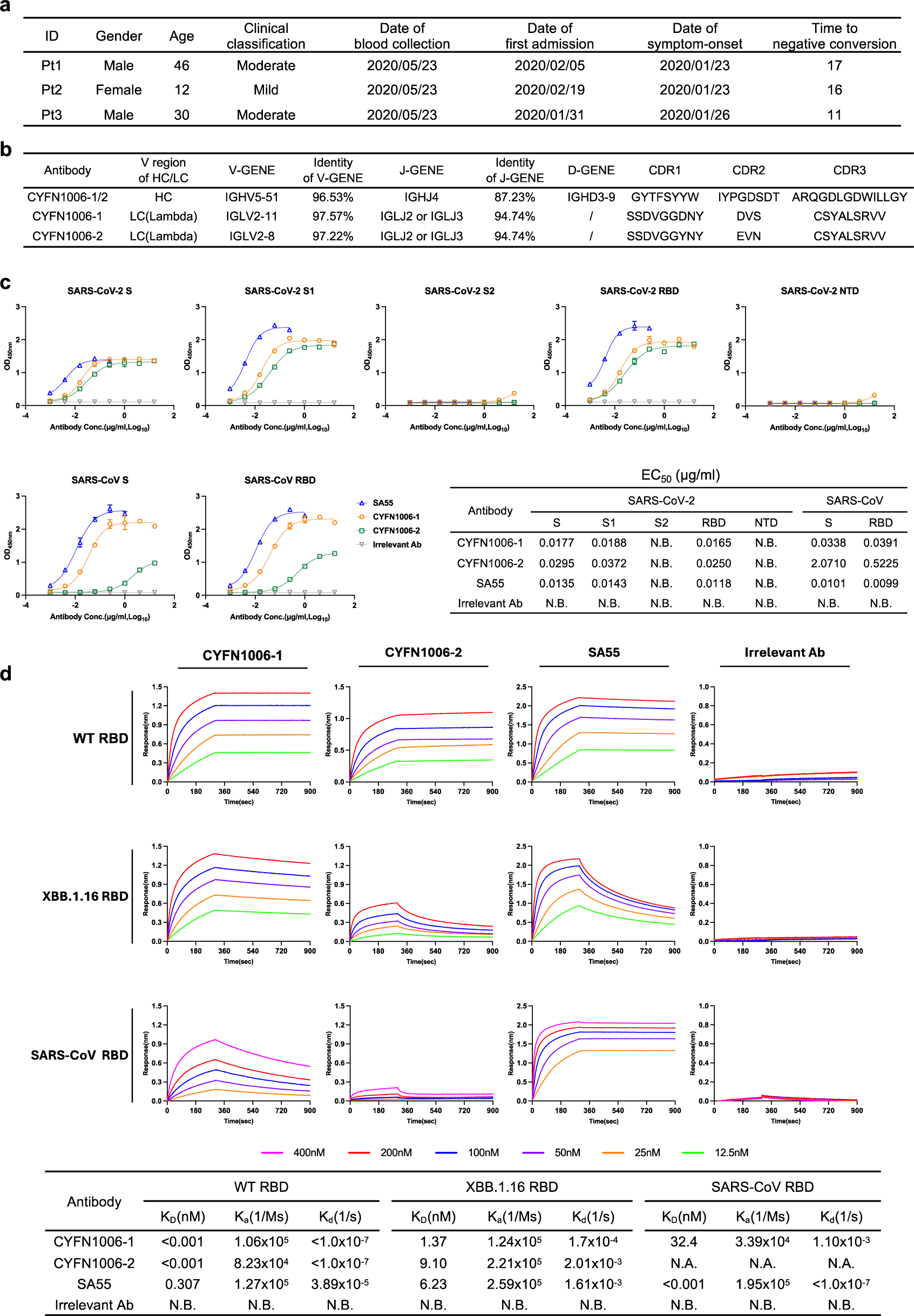
Isolation of antibodies with cross-activity against SARS-CoV-2 and SARS-CoV. **a,** The clinical profiles of the three SARS-CoV-2 convalescent donors. **b,**The gene usage and CDR sequences of the cloned mAbs exhibiting cross-activity against both SARS-CoV-2 and SARS-CoV. **c,** Binding of the indicated mAbs to S, S1, S2, RBD, NTD of SARS-CoV-2 and S, RBD of SARS-CoV, as measured by ELISA. **d,** Binding affinity of the indicated mAbs to SARS-CoV-2 WT RBD, XBB.1.16 RBD and SARS-CoV RBD was analyzed by BLI. N.B., no binding. N.A., not applicable, binding too weak to accurately calculate.

We assessed the binding activity of these two mAbs to SARS-CoV-2 and SARS-CoV, in comparison with SA55, a broad sarbecovirus neutralizing antibody isolated from SARS-CoV-2-vaccinated SARS convalescents^20^. An irrelevant antibody (3G2, a ZIKV NS1-specific Ab^28^) was included as a negative control. Both CYFN1006-1 and CYFN1006-2 demonstrated strong binding to the S1 subunit of the SARS-CoV-2 S trimer, while showing no binding to the S2 subunit, as detected by ELISA assay. Within the S1 subunit, both mAbs bound to the RBD, but not to the NTD. Although CYFN1006-1 and CYFN1006-2 exhibited similar binding to SARS-CoV-2, their binding patterns to SARS-CoV S and RBD appeared different. Specifically, CYFN1006-1 showed robust binding to SARS-CoV S and RBD, whereas CYFN1006-2 displayed significantly weaker binding to the SARS-CoV antigens (**Fig. 1c**).

Their binding affinity to the RBDs of SARS-CoV-2 WT, XBB.1.16, and SARS-CoV was further quantified using Bio-Layer Interferometry (BLI), as illustrated in **Fig. 1d**. CYFN1006-1 exhibited very strong binding (<0.001 nM) to SARS-CoV-2 WT RBD, reduced but still substantial (1.37 nM) binding to SARS-CoV-2 XBB.1.6 RBD, and further reduced (32.4 nM) binding to SARS-CoV RBD. Meanwhile, CYFN1006-2 also demonstrated very strong binding to SARS-CoV-2 WT RBD and reduced binding to XBB.1.6 RBD. However, its binding affinity to SARS-CoV RBD was diminished to a level too low to accurately measure. In contrast, SA55 exhibited the highest binding affinity to SARS-CoV RBD but reduced affinity to SARS-CoV-2 RBDs.

### Potent and broad neutralizing activity of CYFN1006-1 and CYFN1006-2

Next, we sought to evaluate the breadth of neutralization achieved by these two antibodies. We compiled a comprehensive panel of 42 pseudoviruses, encompassing all representative SARS-CoV-2 variants, SARS-CoV, bat or pangolin coronaviruses in the SARS-CoV-2-related lineage (RaTG13, GD-Pangolin) and SARS-CoV-related lineage (WIV1, SHC014), as well as MERS-CoV (**Fig. 2a and Extended Data Fig. 2**). The SARS-CoV-2 variants in our panel included previously identified Variants of Concern (VOCs) (B.1.1.7, B.1.351, P.1, B.1.617.2, BA.1) and Variants of Interest (VOIs) (B.1.525, B.1.621, C.37, etc), comprising variants that were once prevalent such as BA.2, BA.2.12.1, BA.2.75, BA.4/5, BQ.1.1, CH.1.1, and various XBB subvariants, as well as those recently identified and currently circulating variants such as JN.1 and KP.2. The mutations dispersed across the virus S protein are depicted in **Fig. 2b**.

**Fig. 2:**
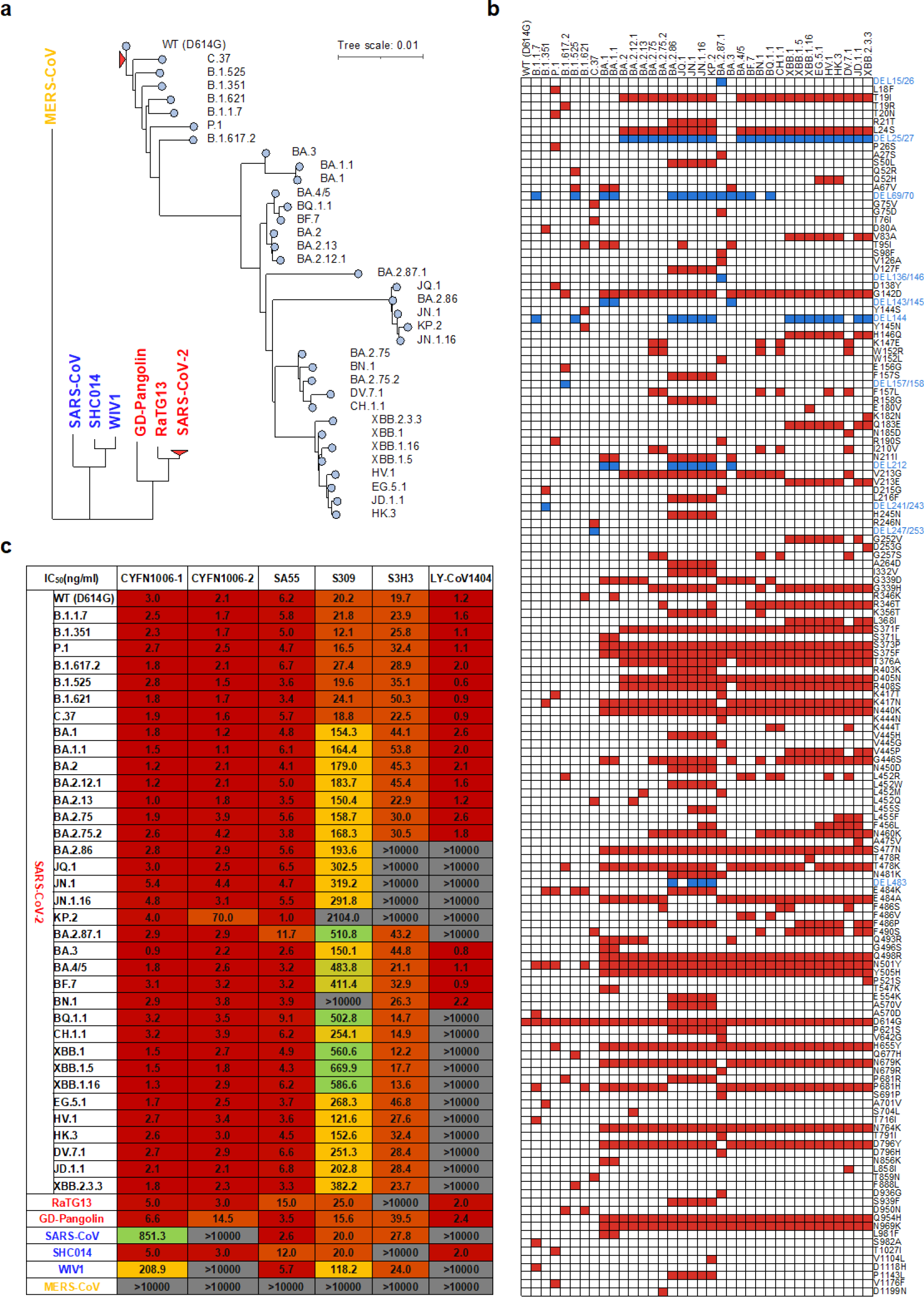
Broad-spectrum neutralizing activity of the mAbs against SARS-CoV-2 variants, SARS-CoV, and other sarbecoviruses. **a,** Phylogenetic tree of the tested comprehensive panel of pseudotyped viruses, including SARS-CoV-2 variants, SARS-CoV, and their related sarbecoviruses, as well as MERS-CoV. **b,** Matrix of mutations on S protein across SARS-CoV-2 variants. **c**, Neutralization IC values of the six_50_ indicated mAbs against the comprehensive panel of pseudotyped viruses.

We assessed the neutralization potency of CYFN1006-1 and CYFN1006-2 on this panel of pseudoviruses, with four previously reported mAbs (SA55, S309, S3H3, and LY-CoV1404) for comparison. As previously reported, LY-CoV1404 completely lost its activity against descendants of BQ, CH, XBB, as well as the JN.1 and KP.2 variants. The SD1-targeting mAb S3H3 was rendered completely inactive against the BA.2.86 and JN.1 variants, likely due to the presence of the E554K mutation in the antibody’s epitope. S309 retained activity, albeit significantly reduced, against most post-Omicron variants, but it completely lost activity against BN.1 and KP.2 variants, likely due to the combinatorial effects of the R346T and K356T mutations (**Fig. 2b-c**).

In contrast, CYFN1006-1 demonstrated consistent neutralization of all tested SARS-CoV-2 variants with unaffected potency, displaying IC_50_ values ranging from approximately 1 to 5 ng/mL, comparable to or even superior to those of SA55. Similarly, CYFN1006-2 exhibited high potency against all SARS-CoV-2 variants, with a slightly reduced efficacy against KP.2. In terms of their activity against the extended sarbecoviruses, both CYFN1006-1 and CYFN1006-2 efficiently neutralized SARS-CoV-2-related viruses (RaTG13, GD-Pangolin). However, only CYFN1006-1 maintained reduced neutralization potency against SARS-CoV and the SARS-CoV-related lineage (WIV1, SHC014), while CYFN1006-2 did not neutralize SARS-CoV or WIV1. As anticipated, none of these monoclonal antibodies neutralized MERS-CoV (**Fig. 2c**).

Given the outstanding neutralization activity of CYFN1006-1 and CYFN1006-2 against pseudoviruses, we extended our testing to authentic viruses, including various SARS-CoV-2 variants (WT, BA.1, XBB.1, EG.5.1, BA.2.86) and SARS-CoV (GZ50). Consistent with the pseudovirus results, both mAbs effectively neutralized all tested SARS-CoV-2 variants with similarly high potency. For SARS-CoV, CYFN1006-1 demonstrated the ability to completely neutralize the virus, albeit with less potency, whereas CYFN1006-2 did not exhibit neutralizing activity against it (**Extended Data Fig. 3**). Taken together, these results confirm that a single ancestral SARS-CoV-2 infection could elicit antibody response against conserved epitopes, albeit rare, with broad-spectrum neutralization activity against evolving viral variants that the donor had never been exposed to.

### Cryo-EM structure of CYFN1006-1 complexed with SARS-CoV-2 XBB.1.16 S trimer

We therefore next explored the structural basis of the broad neutralization activity of CYFN1006-1 and CYFN1006-2 and tried to explain the mechanism of their difference in neutralization. First, structure of CYFN1006-1 immunoglobulin G (IgG) in complex with the SARS-CoV-2 XBB.1.16 S was determined using cryo-electron microscopy (cryo-EM). The XBB.1.16 S trimer was mixed with CYFN1006-1 at a molar ratio of 1:1.2, incubated at 4℃ for 30 min, purified by gel filtration chromatography. The XBB.1.16 S-CYFN1006-1 complex was used for cryo-EM study (**Extended Data Fig. 4**).

Cryo-EM analysis has unveiled three distinct conformational states of the XBB.1.16 S-CYFN1006-1 complex (**Fig. 3a and Extended Data Fig. 5**). In state 1, two S trimers featuring ‘3 down-RBDs’ each interact with three CYFN1006-1 molecules, resulting in the formation of a di-trimer (5.41 Å). In state 2, XBB.1.16 S trimers with ‘3 down-RBDs’ engage with three CYFN1006-1s (2.87 Å). In state 3, XBB.1.16 S trimers with ‘2 down-RBDs’ bind with three CYFN1006-1s (2.84 Å). The S RBD with CYFN1006-1 Fab region was locally refined to 2.92 Å. The Fc region is not visible in the final structure due to its flexibility.

**Fig. 3:**
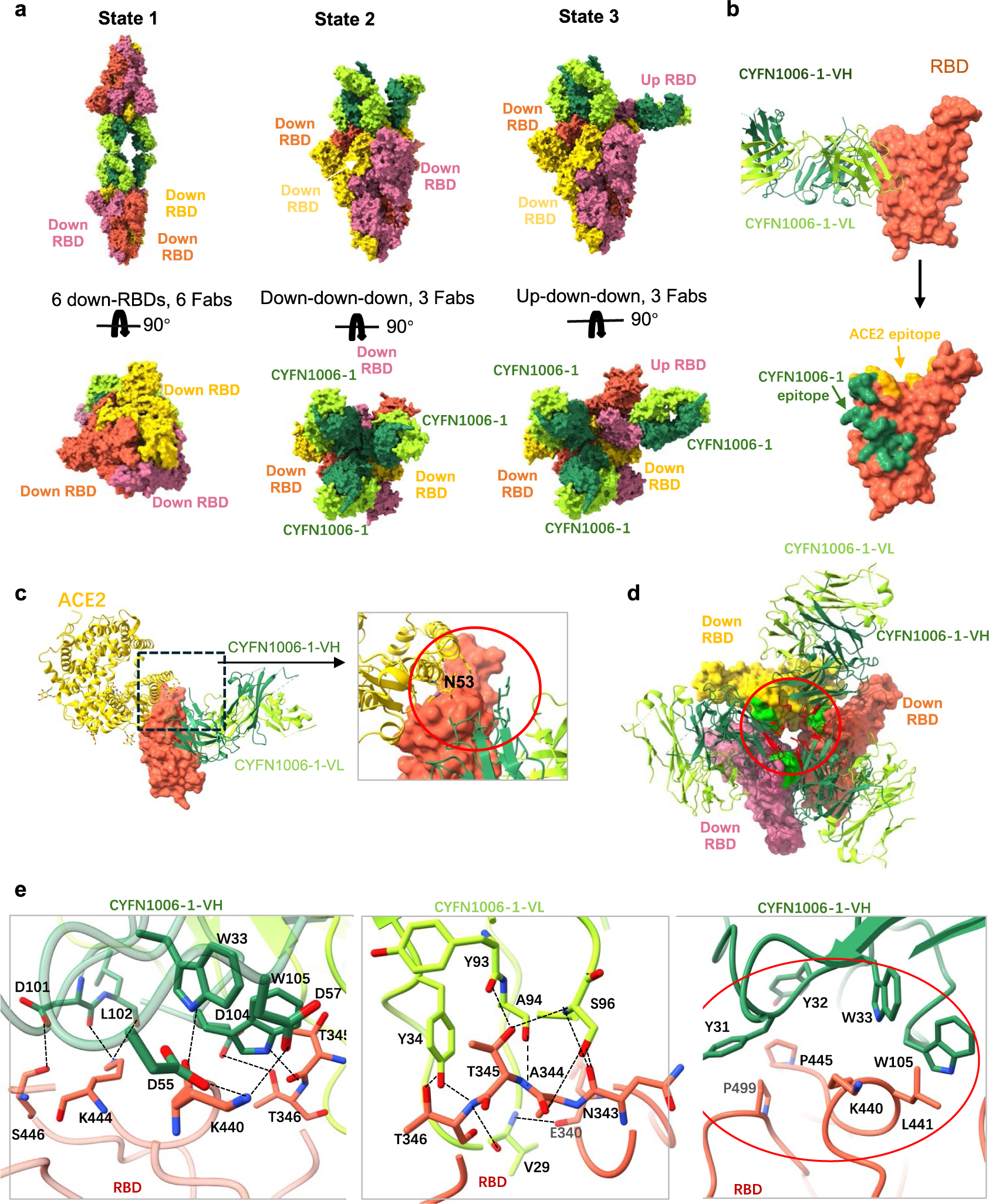
Cryo-EM structure of Omicron XBB.1.16 S trimer in complex with CYFN1006-1. **a,** Cryo-EM structures of the XBB.1.16 S trimer in complex with CYFN1006-1 IgG, shown in surface. **b,** Structure of XBB.1.16 RBD-CYFN1006-1. The RBD is displayed in orange surface mode. The heavy chain and light chain of CYFN1006-1 are shown as ribbons colored in sea green and green yellow, respectively. ACE2 binding epitopes are shown in yellow. **c**, Superimposition of RBD-CYFN1006-1 with RBD-ACE2 (PDB: 7XAZ) reveals the relative position of ACE2 and CYFN1006-1. **d**, Top view of S trimer complexed with CYFN1006-1s, showing that RBDs interacts with the neighboring CYFN1006-1. The RBD and CYFN1006-1 residues involved in the interactions are colored green and red, respectively. **e**, Detailed interactions between the CYFN1006-1 and XBB.1.16 RBD. Hydrogen bonds are represented by dashed lines. Hydrophobic interactions are marked with red circles.

CYFN1006-1 binds to the outer side of RBD, burying a surface area of 933 Å^2^. The epitopes are positioned next to the receptor-binding motif (RBM) of ACE2 (**Fig. 3b**). Structural superposition of the RBD-ACE2 (PDB: 7XAZ) and XBB.1.16 RBD-CYFN1006-1 complex indicates that CYFN1006-1 is close to the glycan on ACE2^N53^ but does not produce steric hindrance conflict with ACE2 (Fig.3c). This is consistent with the ACE2 binding test, which showed that CYFN1006-1 did not prevent the binding between ACE2 and RBD (**Extended Data Fig. 6**). The CYFN1006-1 Fabs on the S trimer can interact with the adjacent Fabs through hydrogen bonding and van der Waals forces between heavy chains (**Extended Data Fig. 7a**). In addition, D62 and S63 of CYFN1006-1 heavy chain can form hydrogen bonds with T500 of the adjacent RBD (**Extended Data Fig. 7b**), locking the RBD in the “down” conformation. Five of six complementary determining regions of CYFN1006-1, are engaged in RBD binding, involving a total of 17 RBD residues (**Fig. 3d and Extended Data Fig. 8**). A rich interaction network was formed, including 18 pairs of hydrogen bonds and 1 hydrophobic interaction patch. V30 of CDRL1 can interact with E340, V341, F347, N354 and K356 of RBD. Y107, A108, S114 of CDRL3 and D112A, W112 of CDRH3 can form 7 pairs of hydrogen bonds with RBD residues N343, A344, T345. The D62 and D64 of CDRH2 form two pairs of hydrogen bonds with the K440 of RBD. D109-W112 of CDRH3 can form 5 pairs of hydrogen bonds with S446, K444, T346. Notably, N343, A344, and T345 play an important role in stabilizing CYFN1006-1 binding to RBD, and are highly conserved in various mutant strains, which are important sites for antibody drug development and vaccine design. K440, L441, P445, and P499 of RBD are embedded in the hydrophobic patch formed by CDRH1 (Y36, Y37, W38) and CDRH3 (W112) to further stabilize the CYFN1006-1 binding to RBD (**Fig. 3e**). In addition, S399 and R509 of RBD are also involved in CYFN1006-1 binding through van der Waals forces.

In summary, CYF1006-1 may neutralize SARS-CoV-2 by locking RBD in a “down” conformation to prevent S trimer from binding to ACE2, as well as by cross-linking S trimers to become di-trimers to block virus infection.

### Cryo-EM structure of CYFN1006-2 complexed with SARS-CoV-2 EG.5.1 S trimer

In addition, we used cryo-EM to analyze the structure of EG.5.1 S-CYFN1006-2 complex using the same method as XBB.1.16 S-CYFN1006-1 complex. Cryo-EM analysis revealed only one conformation of EG.5.1 S-CYFN1006-2 complex. That is, two S trimers with “3 down-RBDs” bind three CYFN1006-2 molecules, forming a di-trimer (**Fig. 4a**), similar to CYF1006-1 state1. One S trimer with three CYFN1006-2 Fab is locally refined to 3.04 Å, and the RBD-CYFN1006-2 Fab region was locally refined to 2.96 Å (**Extended Data Fig. 9**).

**Fig. 4:**
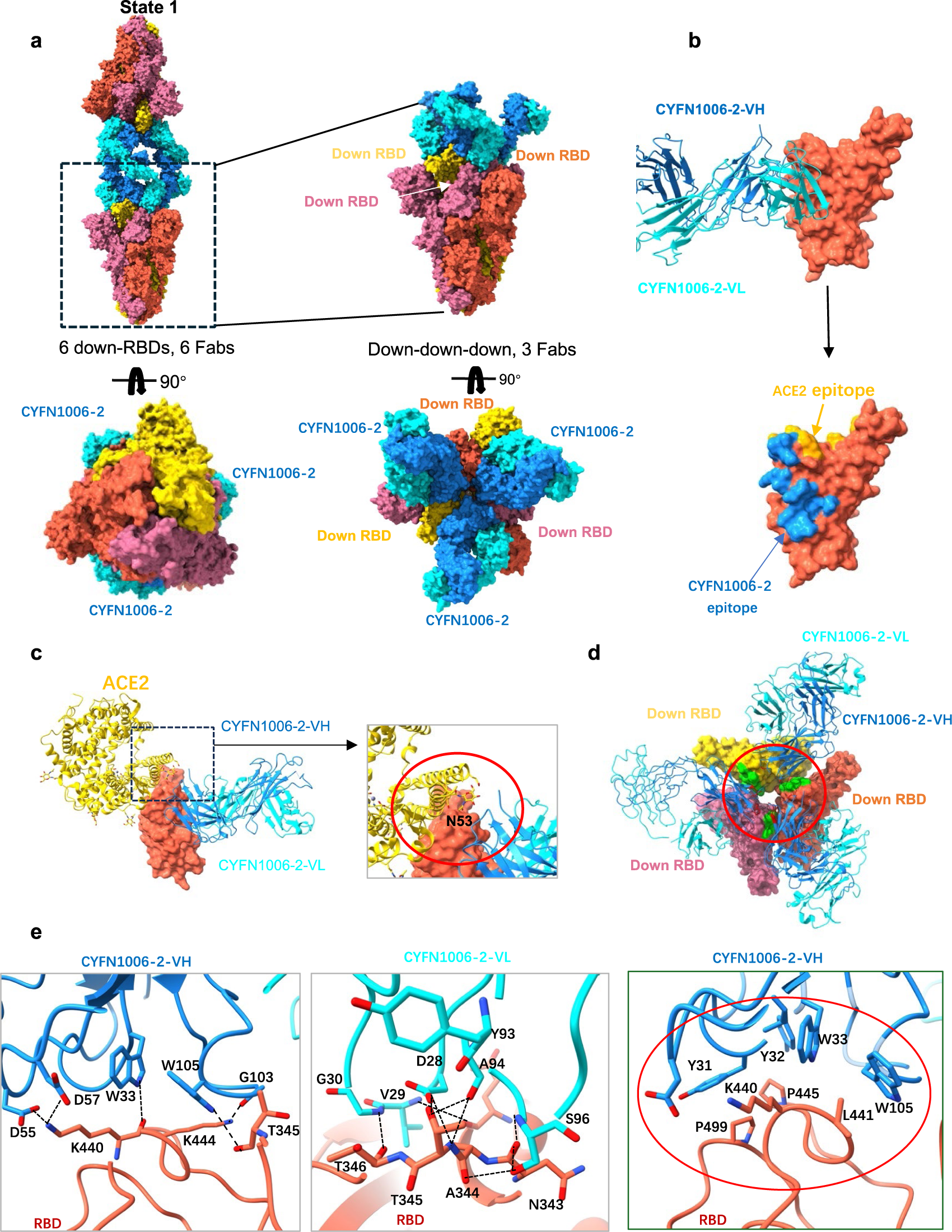
Cryo-EM structure Omicron EG.5.1 S trimer in complex with CYFN1006-2. **a**, Cryo-EM structures of the EG.5.1 S trimer in complex with the CYFN1006-2 IgG, shown in surface. **b**, Structure of EG.5.1 RBD-CYFN1006-2. The RBD is displayed in orange surface mode. The heavy chain and light chain of CYFN1006-2 are shown as ribbons colored in dodger blue and cyan, respectively. ACE2 binding epitopes are shown in yellow. **c**, Superimposition of RBD-CYFN1006-2 with RBD-ACE2 (PDB: 7XAZ) revealed the relative position of ACE2 and CYFN1006-2. **d**, Top view of S trimer complexed with CYFN1006-2s, showing that RBDs interact with the neighboring CYFN1006-2. The RBD and CYFN1006-2 residues involved in the interactions are colored green and red, respectively. **e**, Detailed interactions between the CYFN1006-2 and EG.5.1 RBD. Hydrogen bonds are represented by dashed lines. Hydrophobic interactions are marked with red circles.

Structural analysis shows that the epitopes of CYFN1006-2 with RBD are very similar to those of CYFN1006-1. When compared to CYFN1006-1, the epitopes of CYFN1006-2 are slightly shifted away from RBM and do not interact with R509 next to RBM. The binding surface area is 875 Å^2^, involving a total of 16 amino acids, forming 14 pairs of hydrogen bonds and a hydrophobic plaque (**Fig. 4b**). Structural superposition of the RBD-ACE2 (PDB: 7XAZ) and EG.5.1 RBD-CYFN1006-2 complex indicates that the CYFN1006-2 is adjacent to the glycosylated residue ACE2^N53^, but does not clash with ACE2 (**Fig. 4c**). This is also consistent with the ACE2 binding test (**Extended Data Fig. 6**). Three CYFN1006-2 Fabs bound to the same S trimer with the heavy chains interacting with each other by van der Waals forces. In addition, the binding mode of CYFN1006-2 is similar to that of CYFN1006-1, S63 of each heavy chain can form hydrogen bonds with T500 of the adjacent RBD, preventing the RBD from forming “up” conformation (**Fig. 4d** and **Extended Data Fig. 7c**). K440, L441, P445 and P499 of RBD are embedded in the hydrophobic bags of CDRH1 (Y36, Y37, W38) and CDRH3 (W112) to further stabilize the combination of CYFN1006-2 and RBD. N343, N344, T345 of RBD are highly conserved in a variety of SARS-CoV-2 mutants, and can form 8 pairs of hydrogen bonds with D29 of CDRL1, Y107, A108, S114 of CDRL3, and W112 of CDRH3, which may be the reason for the broad-spectrum neutralization of CYFN1006-2. K440, K444, and T346 participate in the formation of 5 pairs of hydrogen bonds (**Fig. 4e**). In addition, RBD residues H339, E340, V341, F347, N354, S399, and S446 are also involved in CYFN1006-2 binding through van der Waals forces.

Therefore, CYF1006-2 can neutralize SARS-CoV-2 by similar mechanisms as CYF1006-1, such as maintaining the RBD in the down conformation, crosslinking S trimers to di-trimers. However, what makes the difference between CYF1006-1 and CYF1006-2 in neutralizing SARS-CoV? Since the binding affinity of CYF1006-2 to SARS-CoV RBD is too low, we superimposed SARS-CoV RBD (PDB: 2DD8) with SARS-CoV-2 RBD to predict the interactions between SARS-CoV RBD and CYFN1006-1/CYFN1006-2. Structural analysis revealed that CYFN1006-1 may form three more pairs of hydrogen bonds with SARS-CoV RBD than CYFN1006-2 (**Extended Data Fig. 10a**). This finding is consistent with and can explain the binding and neutralization results, which demonstrated that CYFN1006-1 bound to and neutralized SARS-CoV much more effectively than CYFN1006-2. Furthermore, our results also showed that the main difference of the neutralizing activity of CYFN1006-1 and CYFN1006-2 among the SARS-CoV-2 variants was against the KP.2 variant, with CYFN1006-2 showing significant weaker activity than that of CYFN1006-1. We predicted KP.2 RBD-antibody complex models, showing that CYFN1006-1 could form more hydrogen bonds with KP.2 RBD than CYFN1006-2 (**Extended Data Fig. 10b**). This may explain the better neutralization effect of CYFN1006-1 on KP.2 than CYFN1006-2.

### Protection of CYFN1006-1 against SARS-CoV-2 Omicron JN.1 in hamsters

The in vitro neutralization results indicated that, compared to CYFN1006-2, CYFN1006-1 exhibited similar activity against all the tested SARS-CoV-2 variants but demonstrated a broader neutralization breadth to cover SARS-CoV and related sarbecoviruses. Therefore, we selected CYFN1006-1 to assess its anti-SARS-CoV-2 activity in vivo using a hamster model. Male hamsters (aged 6-8 weeks) were intranasally challenged with SARS-CoV-2 Omicron JN.1 at 8 hours after being treated with one dose of CYFN1006-1 antibodies (20 mg/kg) through intraperitoneal injection. At day 4 post-virus inoculation, the hamsters were sacrificed to harvest nasal turbinate (NT) and lung tissues for virological assessments (**Fig. 5a**). Our results demonstrated that passive immunization with CYFN1006-1 significantly reduced the replication of JN.1 in the hamster lungs, as evidenced by the significantly lowered viral subgenomic E (sgE) gene copy and infectious virus titers (**Fig. 5b**). In parallel, we performed immunofluorescence staining and histopathological analysis of SARS-CoV-2 nucleocapsid (N) protein in the harvested hamster lungs. Compared with the PBS-treated group, the expression of viral N protein was substantially reduced in the lungs of infected hamsters upon CYFN1006-1 treatment (**Fig. 5c**). Importantly, histopathological examination of the lung showed multi-focal inflammation, collapse and congestion of the alveoli wall, epithelial damage in the small bronchioles, and alveolar collapse in the PBS-treated hamsters. In contrast, these histological changes were markedly attenuated in the lungs of CYFN1006-1-treated hamsters (**Fig. 5d**). Overall, our results reveal that CYFN1006-1 have the potential to neutralize SARS-CoV-2 Omicron JN.1 in vivo.

**Fig. 5:**
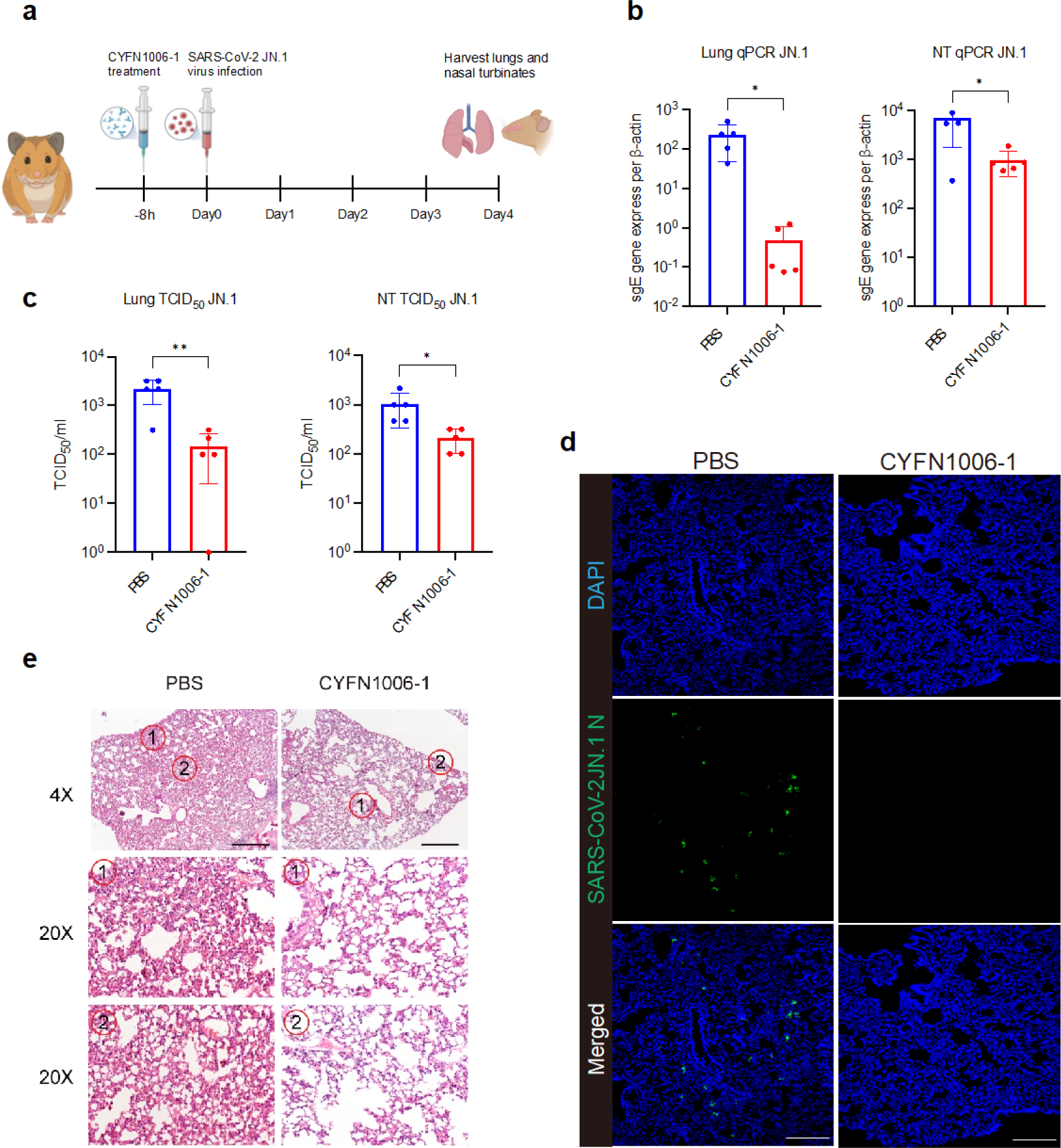
Prophylactic efficacy of CYFN1006-1 against SARS-CoV-2 JN.1 in hamsters. **a,** Schematic of the in vivo mAb efficacy test experiments in hamsters. 6-to-8-week-old male Syrian golden hamsters were treated with one dose of mAb CYFN1006-1 antibody (20 mg/kg, n=5) through intraperitoneal injection antibodies at 8 hours before challenging with 1×10^4^ PFU JN.1. Animals were euthanized at 4 dpi for collection of nasal turbinate (NT) and lung tissues for detection of viral burden and viral antigen expression. **b,c,** Hamster NT and lungs viral sgE copies and infectious titers were measured with (**b**) probe-specific RT-qPCR and (**c**) TCID_50_ assay. **d,** SARS-CoV-2-N protein in hamster lung tissue was stained with anti-SARS-CoV-2-N antibody and visualized using fluorescent microscopy, with DAPI staining used for cell nuclei. Scale bar, 200 μm. **e,** Histological examination of hamster lungs with hematoxylin and eosin (H&E) staining. Scale bar, 200 μm. Data represented means ± SEM from the indicated number of biological repeats. p values were determined by unpaired t test. ^∗^p < 0.05; ^∗∗^p < 0.01.

## Discussion

Neutralizing antibodies with robust autologous neutralization activity often exhibit reduced effectiveness against future variants, especially those obtained early in the pandemic from individuals infected or vaccinated with the ancestral strain. To date, the majority of antibodies exhibiting cross-neutralization activities have been isolated from donors who have experienced prior SARS infection, or multiple vaccinations or infections, or possess hybrid immunity resulting from both vaccinations and infections^20, 27^. In this study, we have successfully identified antibodies capable of potently neutralizing all major SARS-CoV-2 variants, originating from a human donor with ancestral virus exposure exclusively. Notably, considering the timing of sample collection and the donor’s age, it is highly unlikely that this individual had prior exposure to either the SARS-CoV-2 vaccine or SARS-CoV infection. From this donor, we cloned two antibodies, both bearing the rare IGHV5-51 heavy chain gene but paired with different light chain genes. Interestingly, while these antibodies demonstrated similarly high potency in neutralizing SARS-CoV-2 variants, they exhibited varying degrees of efficacy in neutralizing SARS-CoV and related animal sabercoviruses.

In our study, we employed an unbiased antibody discovery method^29^, which involved culturing all MBCs without specific bait, followed by screening the culture supernatant for cross-activity against both SARS-CoV-2 and SARS-CoV. Through this approach, we successfully identified two antibodies with cross-neutralization activity from individuals with solely ancestral virus exposure. Our findings highlight the feasibility of discovering rare yet broad-spectrum neutralizing antibodies capable of effectively targeting future circulating strains, even in convalescent patients who were infected with the prototype strain.

We’ve uncovered the structural foundation behind the broad neutralization activity of these antibodies. The binding epitopes of CYFN1006-1 largely overlap with LY-CoV1404^17^, REGN10987^12^, and S309^18^, situated on the outer surface of RBD near the RBM region (**Extended Data Fig. 11**). The epitopes comprise the important residues of F340, N343, T345, R346, K356 in S309 epitope, forming numerous hydrogen bonds and displaying high conservation. The epitope recognized by CYFN1006-1 shares 10 amino acids with CV38-142^30^, which also utilizes the same IGHV5-51 gene as CYFN1006-1. However, the neutralization of CYFN1006-1 against SARS-CoV-2 (IC_50_ < 5 ng/mL) is much more potent than that of CV38-142 (IC_50_ = 3.46 μg/mL). This difference could be explained by the larger epitope region targeted by CYFN1006-1 on the RBD compared to that of CV38-142. In contrast to LY-CoV1404, CYFN1006-1 lacks high-frequency mutation sites such as N439 and N501, rendering its epitopes more conservative. CYFN1006-1 exerts its neutralizing effect not only by hindering RBD-ACE2 binding by locking RBDs in down position, but also by aggregating virions through cross-linking S trimers to form di-trimers similar to S309. When CYFN1006-1 binds to S of “3 down-RBDs “, the heavy chains of the adjacent Fab interact each other and can form hydrogen bonds with the adjacent RBD, locking the RBD in the “down-RBD” and prevent ACE2 binding. The binding epitopes and neutralization mechanism of CYFN1006-2 closely resemble those of CYFN1006-1, albeit with a slight downward shift. This indicates that broadly neutralizing antibodies could target shared hotspot epitopes while contacting residues resistant to mutations, thereby maintaining their efficacy against evolving viral strains.

Notably, both SA55 and CYFN1006-1 may cover all clinical related viral strains evolved so far (Fig 2). CYFN1006-1 shares two amino acids (P445, P499) with the SA55 epitope^20^, leading to spatial conflict and preventing simultaneous binding to the RBD, as confirmed by competition ELISA tests (**Extended Data Fig. 12**). However, SA55 and CYFN1006-1 bind largely different conserve region of RBD ((**Extended Data Fig. 11**). SA55 directly competes RBD binding to ACE2 ^20^, while CYFN1006-1 does not (**Extended Data Fig. 6**). Therefore, a cocktail with combination of SA55 and CYFN1006-1 may still benefit in covering future evolving mutants of SARS-CoV-2 than individual antibodies if used as drugs. A cocktail of three distinct potent and broad neutralizing mAbs may be required to maintain efficacy against evolving viral strains in the future and to reduce the potential risk of emergence of resistant variants associated with the use of single mAb therapies^31, 32^.

Furthermore, in vivo studies in a hamster model demonstrated the protective efficacy of CYFN1006-1 against SARS-CoV-2 infection, providing additional evidence of its therapeutic potential. Overall, these findings emphasize the critical need for continued investment in antibody discovery and development programs to address the ongoing challenges posed by the dynamic nature of the SARS-CoV-2 virus and other related coronaviruses.

## Method

### Peripheral blood mononuclear cell (PBMC) samples

This study was performed with the approval of the Institutional Review Boards (IRB) at the Guangzhou Eighth People’s Hospital (20200134). Blood samples were drawn after each participant provided a written informed consent form. Fresh PBMCs from three COVID-19 patients after four months of SARS-CoV-2 infection were used for B cell isolation using fluorescence-activated cell sorting (FACS). All patients had positive RNA results for SARS-CoV-2 infection and were hospitalized at Guangzhou Eighth People’s Hospital, Guangzhou Medical University, China, in January and February 2020, the early outbreak of COVID-19 in China.

### Expression of S proteins and their subdomain proteins

The proteins for epitope mapping of the isolated antibody were expressed in mammalian cells. Briefly, the coding sequence of the S ectodomain of SARS-CoV-2 (NC_045512.2) or SARS-CoV (NC_004718.3) was inserted into pcDNA3.1 expression vector (Invitrogen, Waltham, MA) with a D7-tag on the C-terminus of each S protein gene as previously reported^29^. Similarly, the expression vectors for S subdomain proteins (S1, S2, RBD, NTD for SARS-CoV-2 and RBD for SARS-CoV) were also constructed. The recombinant plasmid was transfected into mammalian HEK293F cells. After culturing, S protein-containing supernatants were collected and clarified by centrifugation. Aliquots of supernatants were added with a protease inhibitor and then stored at −80℃ until use.

### Isolation and cloning of mAbs

Patient-derived mAbs against SARS-CoV-2 S protein were generated from memory B cells (CD19^+^IgD^-^IgM^-^CD27^+^CD38^low^) in the PBMCs from COVID-19 patients using an *in vitro* B cell stimulation approach as previously reported^33^. Briefly, memory B cells in fresh PBMCs from convalescent COVID-19 patients were sorted into 96-well culture plates with 50 cells per well. B cells were cultured for 7-10 days in culture medium supplemented with growth factors and stimuli including IL-21, Chk2 inhibitor, and CpG2006 in the presence of feeder cells (irradiated PBMCs from healthy blood donors). The culture supernatants were screened for S protein binding using a capture ELISA. The ELISA microplates precoated with anti-D7 tag Ab were used to capture the S protein with D7-tag in the supernatant of HEK293F cells transfected with recombinant pcDNA3.1. Cells in the positive wells were subjected to RNA extraction for cDNA synthesis using the SuperScript™ III First-Strand Synthesis System (Invitrogen, Waltham, MA). The cDNA was used as a template of the nested PCR with gene-specific primers or primer mixes to amplify the transcripts of antibody heavy and light variable genes as previously reported^34^. The PCR products were purified from agarose gel using QIAEX II Gel Extraction Kit (Qiagen, Hilden, Germany). Purified PCR products were digested with restriction enzymes AgeI/SalI, AgeI/BsiWI, or AgeI/XhoI (ThermoFisher Scientific, Waltham, MA) and followed by ligation into human IgG1, Igκ, or Igλ expression vectors, respectively. The recombinant Ab-expressing constructs were sequenced and analyzed using the IMGT/V-QUEST (IMGT, France), a sequence alignment software for the immunoglobulin sequences of the variable regions.

### Epitope mapping

For epitope mapping of the isolated mAbs, a panel of SARS-CoV-2 and SARS-CoV S, and their subdomain proteins (S1, S2, RBD, NTD for SARS-CoV-2 and RBD for SARS-CoV) were used. The binding of each mAb to these proteins was measured using a capture ELISA. Human mAbs starting at 16 µg/ml were used for preparing a 4-fold serial dilution and applied to the capture ELISA. All samples were detected in duplicate. A Zika NS1-specific mAb (the irrelevant mAb) was used as a negative control. The concentration for 50% of maximal effect (EC_50_) was calculated using GraphPad Prism 8.

### Affinity measurement

The K_D_ measurement of isolated mAbs to SARS-CoV-2 S protein were conducted on the Octet K2 System (ForteBio, Fremont, CA) using the Bio-Layer Interferometry (BLI). Briefly, the Streptavidin (SA) sensor (Sartorius,18-5019) captured the biotin-labeled (EZ-Link^TM^ NHS-PEG12-Biotin, ThermoFisher Scientific, A35389) mAbs at 10 µg/ml, then serially diluted RBD protein were added. Three types of RBD proteins were measured for each Ab, including SARS-CoV-2 wild-type, XBB.1.16 (Vazyme, Nanjing, China) and SARS-CoV (ABclonal, Wuhan, China). After washing, the association and dissociation curves were obtained. The K_D_ values were calculated using Data Analysis SPSS 11.0. A Zika NS1-specific mAb (the irrelated mAb) was used as a negative control.

### Antibody competition for binding of ACE2 to RBD and Spike trimer

The competition assay was conducted on the Octet K2 System (ForteBio, Fremont, CA) using the Bio-Layer Interferometry (BLI). To measuring antibody competition for binding of ACE2 to RBD, the Streptavidin (SA) sensor (Sartorius,18-5019) captured the biotin-labeled (EZ-Link^TM^ NHS-PEG12-Biotin, ThermoFisher Scientific, A35389) anti-D7 antibody at 40 µg/ml, which subsequently captured the SARS-CoV-2 RBD protein in the supernatant. After equilibrium, antibody (400nM) and ACE2 protein (400nM) or ACE2 protein and antibody were added in tandem. For control, we also determined the responses using antibody or ACE2 (400 nM) only. To measuring antibody competition for binding of ACE2 to Spike trimer, the ACE2 (200nM) were immobilized to Amine Reactive 2nd-Generation (AR2G) biosensors (Sartorius,18-5092) following the protocol recommended by the manufacturer. Antibody (400nM) plus Spike trimer (100nM) premixed at room temperature for 10 minutes to ensure full reaction. The premixture was then presented to the biosensors loaded with ACE2. The mixture of buffer and Spike trimer was used for a control.

### Production of pseudoviruses

Plasmids encoding the S of SARS-CoV, SARS-CoV-2 and its variants, as well as spikes from SARS-CoV and SARS-CoV-2 related sarbecoviruses, were synthesized. Pseudoviruses were generated by co-transfection of HEK-293T cells with the indicated spike gene using PEI. Cells were cultured overnight at 37 °C with 5% CO_2_ and VSV-G pseudo-typed ΔG-luciferase (G*ΔG-luciferase, Kerafast) was used to infect the cells at a multiplicity of infection of 5 in DMEM with 10% fetal bovine serum (FBS), followed by washing the cells with PBS/2% three times and replaced culture medium by DMEM with 2% FBS. After 24 h, the transfection supernatant was collected and clarified by centrifugation at 4,000 rpm for 10 min. Each viral stock was subsequently incubated with 20% I1 hybridoma (anti-VSV-G; CRL-2700, ATCC) supernatant for 1 hour at 37°C to neutralize the contaminating VSV-G pseudotyped ΔG-luciferase virus before titration and aliquoting for storage at −80°C.

### Pseudovirus neutralization assay

Neutralization assays were performed by incubating pseudoviruses with serial dilutions of monoclonal antibodies and assessing the reduction in luciferase gene expression, as previously described^35^. In brief, Vero-E6 cells were seeded in a 96-well plate at a concentration of 2×10^4^ cells per well. The following day, pseudoviruses were incubated with serial dilutions of the specified monoclonal antibody in triplicate for 30 minutes at 37°C. The mixture was then added to cultured cells and incubated for an additional 24 hours. Subsequently, the luminescence was measured using the Luciferase Assay System (RG062M, Beyotime). IC_50_ was defined as the dilution at which the relative light units were reduced by 50% compared to the virus control wells (virus and cells), after subtraction of the background in the control groups with cells only. IC_50_ values were calculated using nonlinear regression in GraphPad Prism.

### Viruses and biosafety

SARS-CoV GZ50 (GenBank accession number AY304495) was an archived clinical isolate at the Department of Microbiology, HKU. SARS-CoV-2 WT D614G (GISAID: EPL_ISL_497840), BA.1 (GISAID: EPI_ISL_6841980), XBB.1 (GISAID: EPI_ISL_15602393), EG.5.1 (GISAID: EPI_ISL_18461518), BA.2.86 (GISAID: EPI_ISL_18986956) and JN.1 (GISAID: EPL_ISL_18841631) strains were isolated from the respiratory tract specimens of laboratory-confirmed COVID-19 patients in Hong Kong. SARS-CoV-2 was cultured using Vero-E6-TMPRSS2. SARS-CoV was cultured in Vero-E6 cells. All the viruses were titrated by plaque assays. All experiments with infectious SARS-CoV-2 and SARS-CoV were performed according to the approved standard operating procedures of the Biosafety Level 3 facility at the Department of Microbiology, HKU^36^.

### Authentic virus neutralization

An end-point dilution assay in a 96-well plate format was performed to measure the neutralization activity of select purified bsAbs as described previously^27^. In brief, each antibody was serially diluted (fivefold dilutions) starting at 50 μg/mL. Triplicates of each antibody dilution were incubated with indicated live virus at a MOI of 0.1 in DMEM with 7.5% inactivated fetal calf serum for 1 hour at 37 °C. After incubation, the virus–antibody mixture was transferred onto a monolayer of Vero-E6 cells grown overnight. The cells were incubated with the mixture for 70 hours. The cytopathic effects were visually scored for each well in a blinded fashion by two independent observers. The results were then converted into percentage neutralization at a given concentration, and means ± SEM were plotted using a five-parameter dose–response curve in GraphPad Prism.

### Hamster protection and Virus challenge

The use of all animals in the study was approved by The Committee on the Use of Live Animals in Teaching and Research (CULATR) of The University of Hong Kong (HKU). 6-8 weeks old male golden Syrian hamsters were obtained from the Centre for Comparative Medicine Research (CCMR) at HKU. The hamsters were housed in cages equipped with individual ventilation systems, maintained at a humidity level of 65%, and kept at an ambient temperature of 21-23°C with a 12-hour day-night cycle for proper care and management. Group sizes were chosen based on statistical power analysis and our prior experience in examining viral titers in SARS-CoV-2-infected hamsters or mice^37, 38^.

To evaluate the antiviral effects of CYFN1006-1, hamsters were pre-treated for 8 hours intraperitoneally with one dose of PBS or mAb CYFN1006-1 (20 mg/kg) before virus inoculation. All intranasal treatment in hamsters was performed under intraperitoneal ketamine (100 mg/kg) and xylazine (10 mg/kg) anesthesia. Virus-infected hamsters were euthanized at 4 dpi to collect NT and lung tissues for immunofluorescence staining, hematoxylin and eosin (H&E) staining, qRT-PCR, or TCID_50_ assays as we previously described^39^.

### RNA extraction and quantitative reverse transcription polymerase chain reaction (qRT-PCR)

Viral RNA was extracted from homogenized animal tissues using RNeasy Mini Kit (74106, Qiagen). After RNA extraction, qRT-PCR was performed using QuantiNova Probe RT-PCR Kit (208354, Qiagen) or QuantiNova SYBR Green RT-PCR Kit (208154, Qiagen) with the LightCycler 480 Real-Time PCR System (Roche, Basel, Switzerland). The primer and probe sequences are available upon request.

### Histology and Immunofluorescence staining

Infected hamster lungs were fixed by 4% paraformaldehyde for 24 hours. Then samples were washed with 70% ethanol and a TP1020 Leica semienclosed benchtop tissue processor (Leica Biosystems, Buffalo Grove, IL, USA) was applied to embed lung samples in paraffin. Embedded samples were sectioned with a microtome (Thermo Fisher Scientific) and placed on microscope slides for drying at 37°C overnight. Sample slides were dewaxed by washing in serially diluted xylene, ethanol, and double-distilled water. For antigens unmasking, diluted antigen unmasking solution (H-3300, Vector Laboratories) was heated until 85°C, then slides were put in solution and boiled for 90s. Unmasked slides were stained with Sudan black B and blocked with 5% fetal bovine serum for 30 min. The in-house rabbit anti-SARS-CoV-2-N immune serum was applied onto samples and incubated at 4°C overnight. Goat anti-rabbit immunoglobulin G(H+L) fluorescein isothiocyanate (65-6111), which functioned as a secondary antibody was purchased from Thermo Fisher Scientific. ProLong™ Diamond Antifade Mountant with DAPI (P36962, Thermo712 Fisher Scientific) was used to stain the cell nucleus while the slides were mounted. Images were taken with the Olympus BX53 fluorescence microscope (Olympus Life Science, Tokyo, Japan).

For H&E staining, tissue sections were stained with Gill’s haematoxylin and eosin. Images were acquired using the Olympus BX53 light microscope (Olympus Life Science, Japan). Five hamsters were sampled in each group and four to six sections from each animal were used for histology analysis.

### Hamster lung and NT tissue infection virus titration

VeroE6-TMPRSS2 was obtained from the Japanese Collection of Research Bioresources (JCRB) Cell Bank. VeroE6-TMPRSS2 was cultured in Dulbecco’s modified Eagle’s medium (DMEM) (11965092, Gibco, USA) supplemented with 10% fetal bovine serum (FBS), 100 units penicillin,100 ug/ml streptomycin, and 2% G418 (ant-gn-5, InvivoGen, China). To quantify the infectious titers of virus-infected hamster lung and NT tissues, a monolayer of VeroE6-TMPRSS2 cells in a 96-well plate was challenged with 10-fold serial diluted supernatants of homogenized tissues. After 96 hours of incubation, JN.1-induced cytopathic effect (CPE) was checked by visualization and calculated as we previously described^40^.

### Protein expression and purification for Cryo-EM

The constructs used to express stable soluble Omicron XBB.1.16 (6P) S and EG.5.1 (6P) S were derived from our previous study^27^. Plasmid was transfected into HEK293F cells by polyethlenimine, supernatant was harvested after 72 hours, and the S trimer is purified with the Ni-Sepharose resin from cytiva Reagents. The target protein was further purified by Superose 6 increase 10/300 column (GE Healthcare) in 20 mM Tris, 200 mM NaCl, pH 8.0 buffer solution.

### Cryo-EM data collection and processing

The XBB.1.16 spike trimer at 5mg/mL was mixed with 5 mg/mL CYFN1006-1 at the molar ratio of 1:1.2 (S trimer /IgG), incubated at 4℃ for 30min, and the complex was separated on Superose 6 Increase 10/300 GL. The size and purity of the complex proteins were then further assessed by SDS-PAGE and dilute to 0.6 mg/mL in 20 mM Tris, pH 8.0, and 200 mM NaCl. The complex of EG.5.1 S and CYFN1006-2 was prepared by the above method. The 3μL composite was added to a new light-emitting porous amorphous Ni-Ti alloy film supported by 400 mesh gold mesh. The sample was immersed in liquid ethane using Vitrobot IV (FEI, Thermo Fisher Scientific) and frozen for 2-s, −3blot force, 10-s waiting time.

Cryo-EM data for XBB.1.16-CYFN1006-1 and EG.5.1-CYFN1006-2 were collected automatically with EPU software on a 300 kV Titan Krios G4 microscope. The device is equipped with a Falcon 4i and Seletrics X-ray imaging filter (Thermo Fisher) with a slit width set to 20 eV. Collect EER movie stack in super resolution mode, XBB.1.16-CYFN1006-1 nominal magnification of 130,000x, EG.5.1-CYFN1006-2 nominal magnification of 105,000x, corresponding physical pixel sizes of 0.932 Å and 1.19 Å. The doses were 1080 frames and 1737 frames, respectively^41^. The defocusing range is −1.0 to −3.0 μm, and the total dose is about 50^e−^/Å^2^. Data processing is performed via modules on RELION v3.1 or cryoSPARC v4.2.1. All micrographs were imported into cryoSPARC for patch CTF estimation^42^, particle pickup and 2D classification. Then select the good particles and import them into Relion for 3D automatic thinning, 3D thinning and 3D classification. The resolution of the report is based on the gold standard Fourier Shell correlation (FSC) 0.143. The data processing flow is summarized in Extended Data Figs. 5 and 9.

### Model building and refinement

EMready^43^ is used to sharpen maps for model building. SWISS-MODEL^44^ is used to generate initial models. The models were fitted into maps using UCSF Chimera^45^, manually built in COOT^46^, followed by multiple rounds of real space refinement in PHENIX^47^. Model validation using PHENIX. Figures were prepared using UCSF Chimera and UCSF ChimeraX^48^. Statistics of data collection and model refinement are shown in Supplementary Table 1.

## Acknowledgements

We thank the Center of Cryo-Electron Microscopy, Core Facility of Shanghai Medical College, Fudan University for the supports on cryo-EM data collection. This study was supported by funding from the National Key Research and Development Program of China (No. 2023YFC3404000 to X.Z.), National Natural Science Foundation of China (32270142 to P.W.), Shanghai Rising-Star Program (22QA1408800 to P.W.), the Program of Science and Technology Cooperation with Hong Kong, Macao and Taiwan (23410760500 to P.W.), Ministry of Science and Technology of China (2021YFC2302500 to L.S.), R&D Program of Guangzhou Laboratory (SRPG22-003 to L.S.), and Guangzhou Municipal Science and Technology Bureau (202201020528 to L.Y.). Pengfei Wang acknowledges support from Open Research Fund of State Key Laboratory of Genetic Engineering, Fudan University (No. SKLGE-2304) and Xiaomi Young Talents Program. Hin Chu acknowledges support from National Natural Science Foundation of China Excellent Young Scientists Fund (Hong Kong and Macau) (32122001); National Key Research and Development Program of China (2021YFC0866100 and 2023YFC3041600); Collaborative Research Fund (C7103-22G) and Theme-Based Research Scheme (T11-709/21-N), the Research Grants Council of the Hong Kong Special Administrative Region; and Health@InnoHK, Innovation and Technology Commission, the Government of the Hong Kong Special Administrative Region.

## Author contributions

P.W., Y.G., H.C., L.S. and L.Y. conceived and supervised the project. X.X., C.L., X.W., J.L., Y.C, R.Q., X.Z., M.G., S.W., and Y.X. conducted the biological experiments. Y.L. and J.S. performed authentic neutralization assays and animal experiments. Y.W. performed the collection of Cryo-EM data and structure determination. W.M. conducted the bioinformatics analysis. P.W., Y.W., Y.L., X.X., C.L., Y.G., H.C., L.S. and L.Y. analyzed the results and wrote the manuscript. All the authors reviewed, commented, and approved the manuscript.

