## Supplementary file for "Potent and broadly neutralizing antibodies against sarbecoviruses elicited by single ancestral SARS-CoV-2 infection"

**Supplementary Table 1.** **Cryo-EM data collection and refinement statistics.**

| | XBB.1.16-CYFN1006-1 | | | | EG.5.1-CYFN1006-2 | | |
|---|---|---|---|---|---|---|---|
| | ditrimer | trimer-3d | trimer-2d1u | local | ditrimer | trimer-3d | local |
| **Data collection and processing** | | | | | | | |
| Magnification | 130,000 | | | | 105,000 | | |
| Voltage (kV) | 300 | | | | 300 | | |
| Total dose (e–/Å$^2$) | 50 | | | | 50 | | |
| Defocus range (μm) | -1.0 to -3.0 | | | | -1.0 to -3.0 | | |
| Pixel size (Å) | 0.932 | | | | 1.19 | | |
| Symmetry imposed | C1 | | | | C1 | | |
| Final particles (no.) | 38112 | 540576 | 936842 | 936842 | 66958 | 244940 | 244940 |
| Map Resolution | 5.41Å | 2.87Å | 2.84 Å | 2.92Å | 3.83 Å | 3.04 Å | 2.96 Å |
| **R.m.s. deviations** | | | | | | | |
| Bond lengths (Å) | 0.001 | 0.002 | 0.002 | 0.005 | 0.002 | 0.002 | 0.003 |
| Bond angles (°) | 0.401 | 0.424 | 0.538 | 0.627 | 0.527 | 0.546 | 0.546 |
| **Validation** | | | | | | | |
| MolProbity score | 1.48 | 1.68 | 1.76 | 2.03 | 1.67 | 1.92 | 1.99 |
| Clash score | 5.30 | 5.27 | 5.62 | 4.27 | 5.28 | 5.85 | 4.11 |
| Rotamer outlier (%) | 1.44 | 1.89 | 2.23 | 3.78 | 1.15 | 2.54 | 2.88 |
| **Ramachandran plot** | | | | | | | |
| Favored (%) | 97.64 | 96.88 | 96.88 | 94.38 | 95.17 | 95.76 | 92.94 |
| Allowed (%) | 2.36 | 3.07 | 3.07 | 5.27 | 4.79 | 4.17 | 6.73 |
| Disallowed (%) | 0 | 0 | 0 | 0 | 0 | 0 | 0 |
| **EMDB** | EMD-39801 | EMD-39802 | EMD-39803 | EMD-39804 | EMD-39805 | EMD-39806 | EMD-39807 |
| **PDB** | 8Z6Q | 8Z6R | 8Z6S | 8Z6T | 8Z6U | 8Z6W | 8Z6X |

**a**

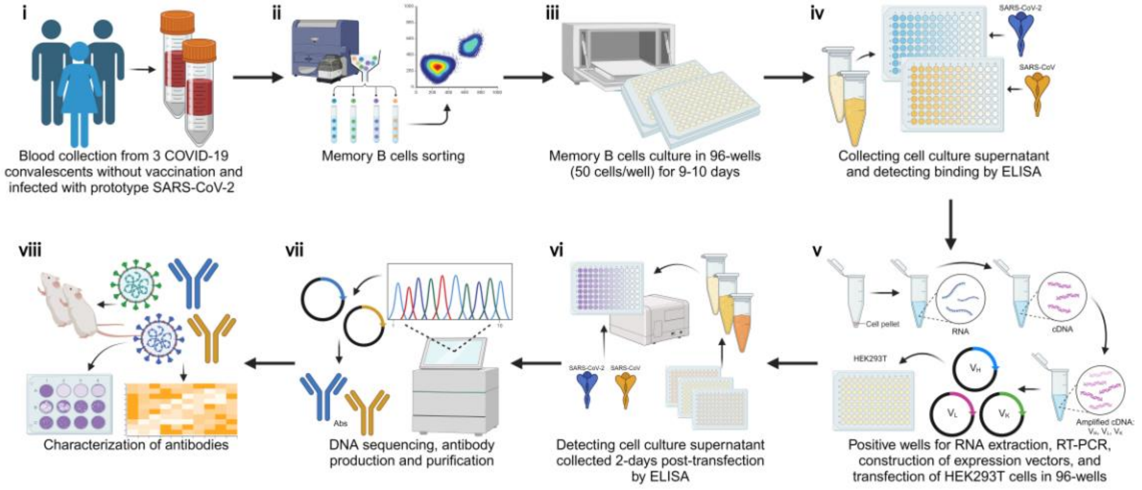


**b**

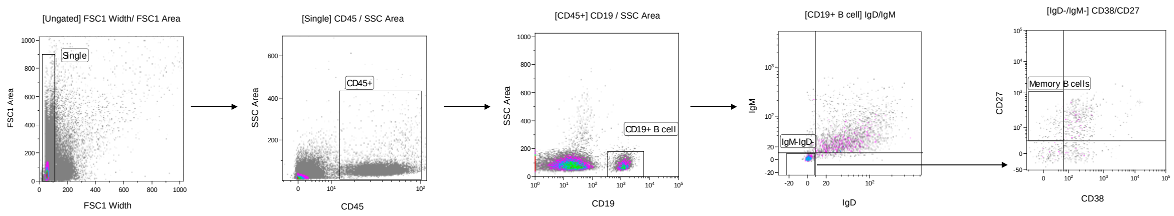


**Extended Data Fig. 1 The process of isolating neutralizing antibodies from COVID-19 convalescents.**

(**a**) Schematic representation of isolating mAbs from 3 COVID-19 convalescent donors. (**b**) The gating strategy for memory B cell Sorting. The first gate was on singlets based on FS TOF vs FS INT. After gating on CD45, then CD19, IgD-IgM- B cells were identified using IgD and IgM expression. IgD-IgM- B cells were further analyzed by CD38 and CD27 expression. $CD19^{+}IgD^{-}IgM^{-}CD27^{+}CD38^{low}$ cells were identified as memory B cells.

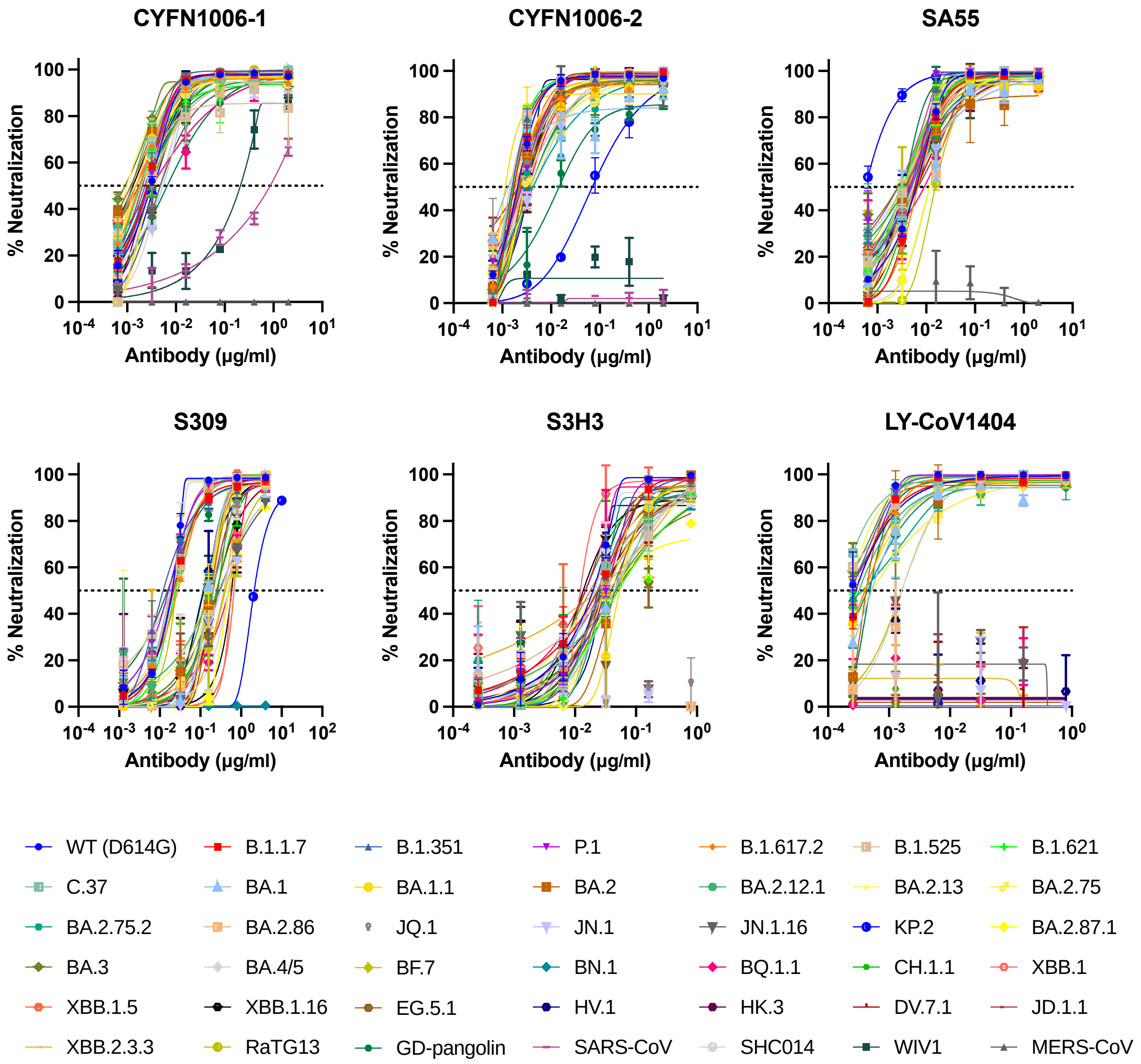


**Extended Data Fig. 2 Neutralization curves of the six mAbs against pseudotyped SARS-CoV-2 variants, SARS-CoV and other related sarbecoviruses.**

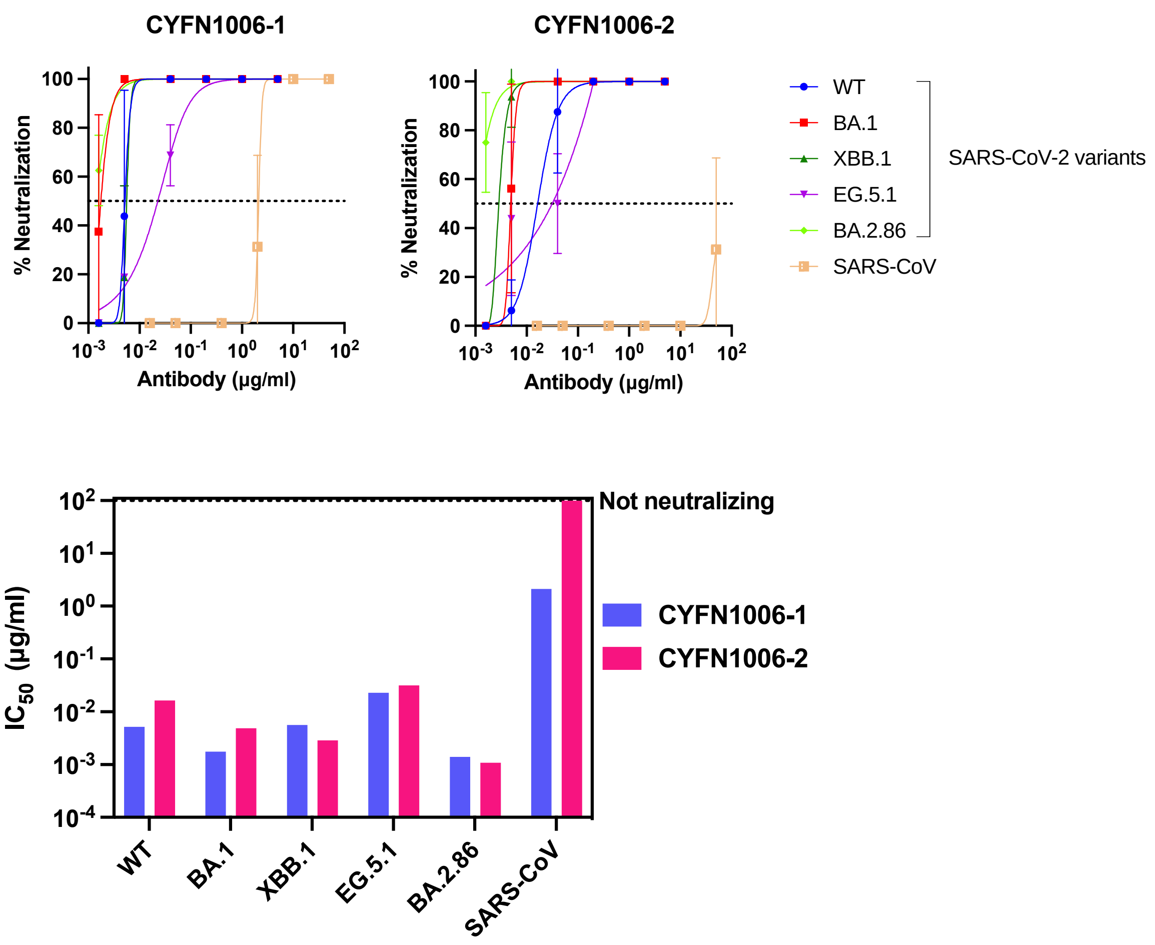


**Extended Data Fig. 3 Authentic virus neutralization.**

(**a**) Neutralization curves of authentic SARS-CoV-2 variants and SARS-CoV by CYFN1006-1 and CYFN1006-2. (**b**) Comparison of $IC_{50}$ values of two indicated mAbs against SARS-CoV-2 variants and SARS-CoV.

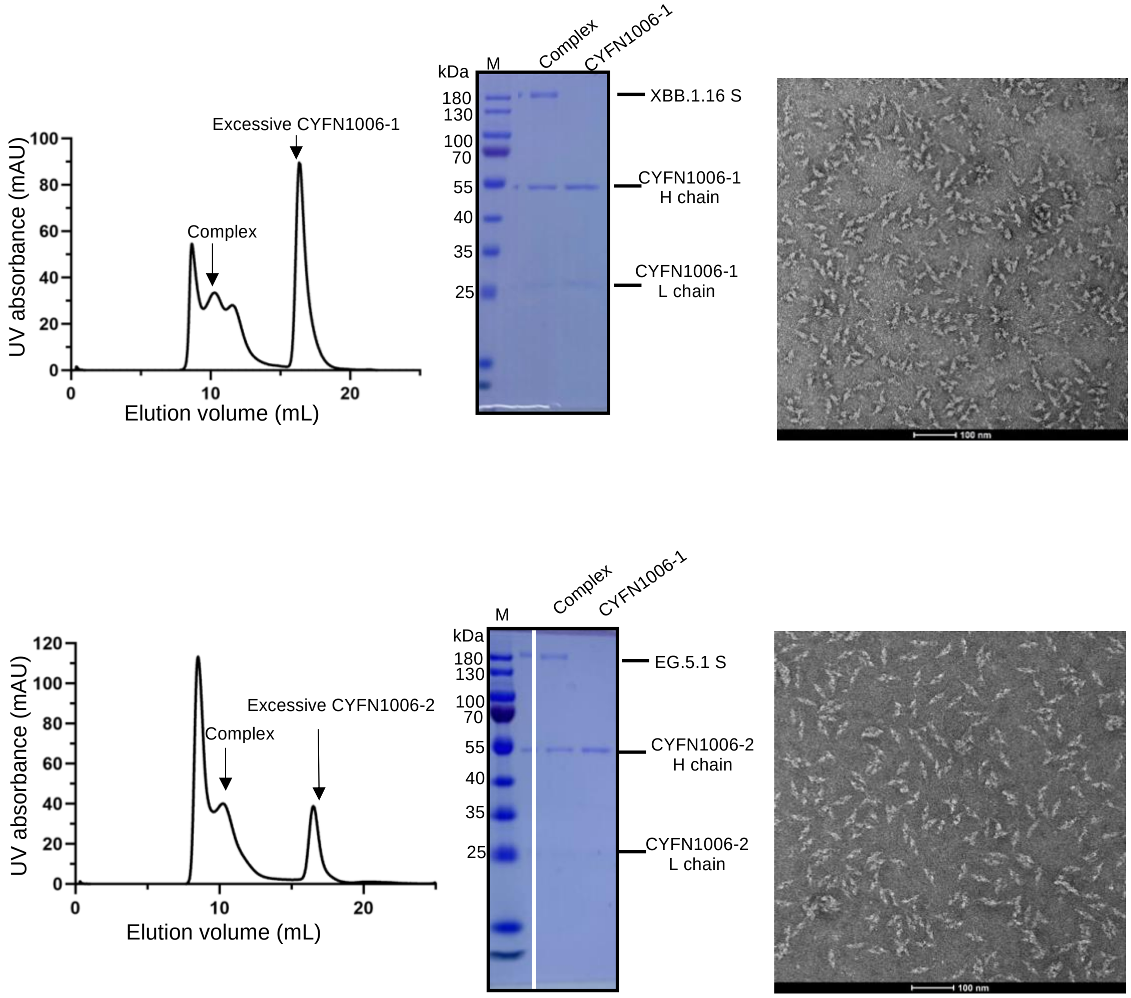


**Extended Data Fig. 4 Sample purification of XBB.1.16 S-CYFN1006-1 and EG.5.1 S-CYFN1006-2 complex.**

(**a**) Purification and negative stain images of XBB.1.16 S-CYFN1006-1 complex. (**b**) Purification and negative stain images of EG.5.1 S-CYFN1006-2 complex.

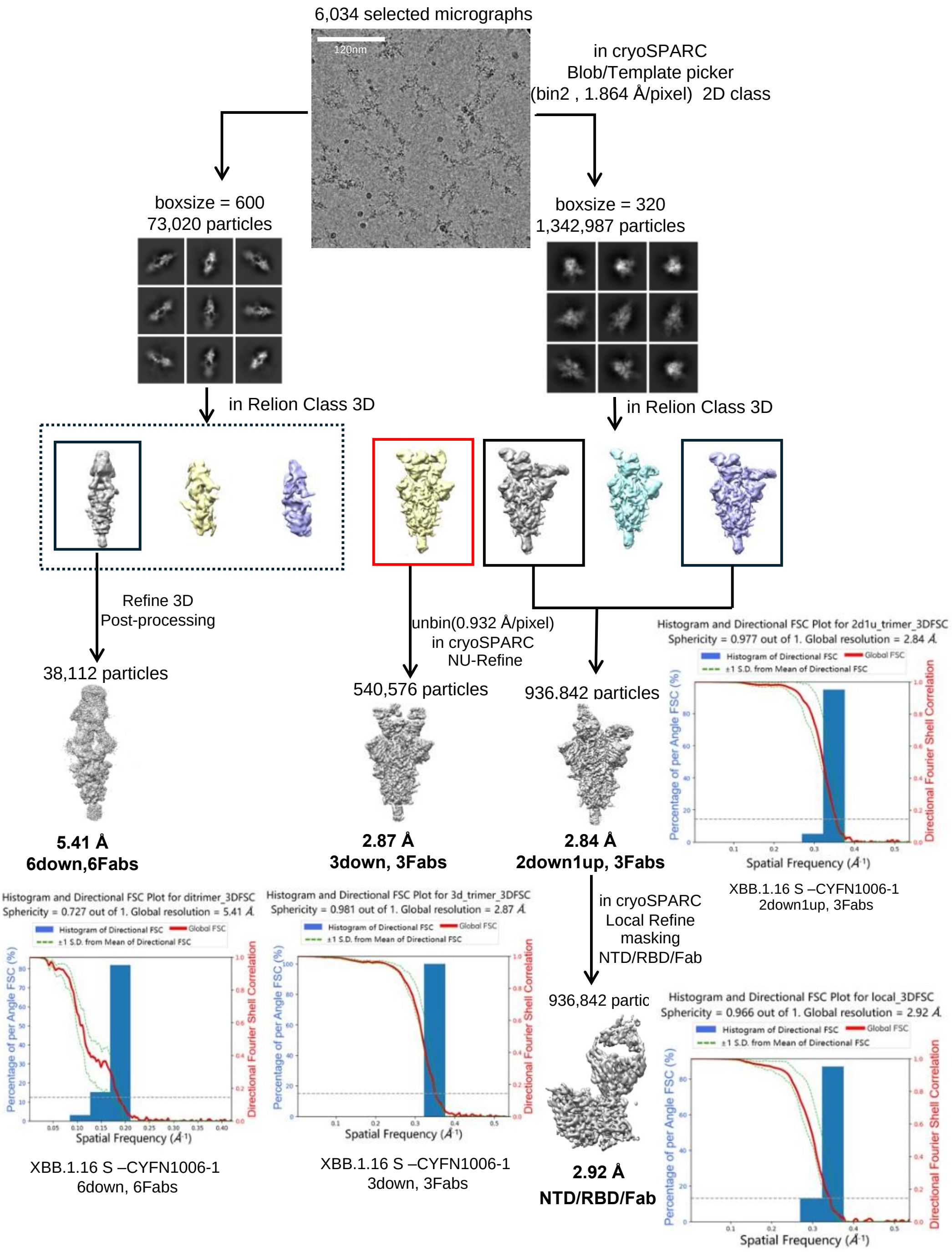


**Extended Data Fig. 5 Data processing flowchart of XBB.1.16 S–CYFN1006-1 complex.**

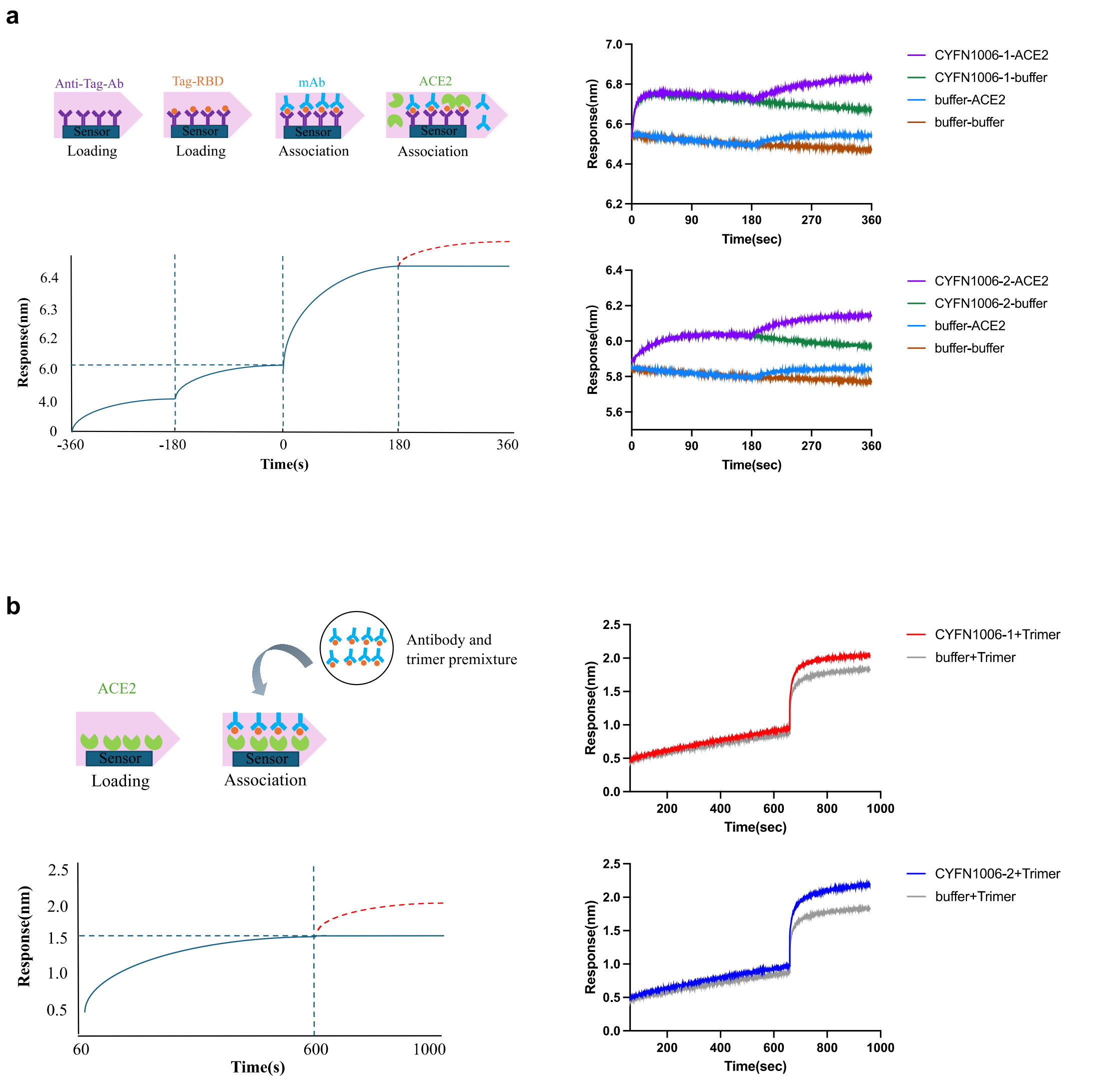


**Extended Data Fig. 6 CYFN1006-1 and CYFN1006-2 did not inhibit the binding of ACE2 to RBD/S as measured by BLI assay.**

(**a**) Binding of ACE2 to RBD was assessed after a first association phase with mAb (CYFN1006-1, CYFN1006-2) or with buffer control. (**b**) Binding of S trimer to ACE2 was assessed with S trimer alone or with premixed S/mAb.

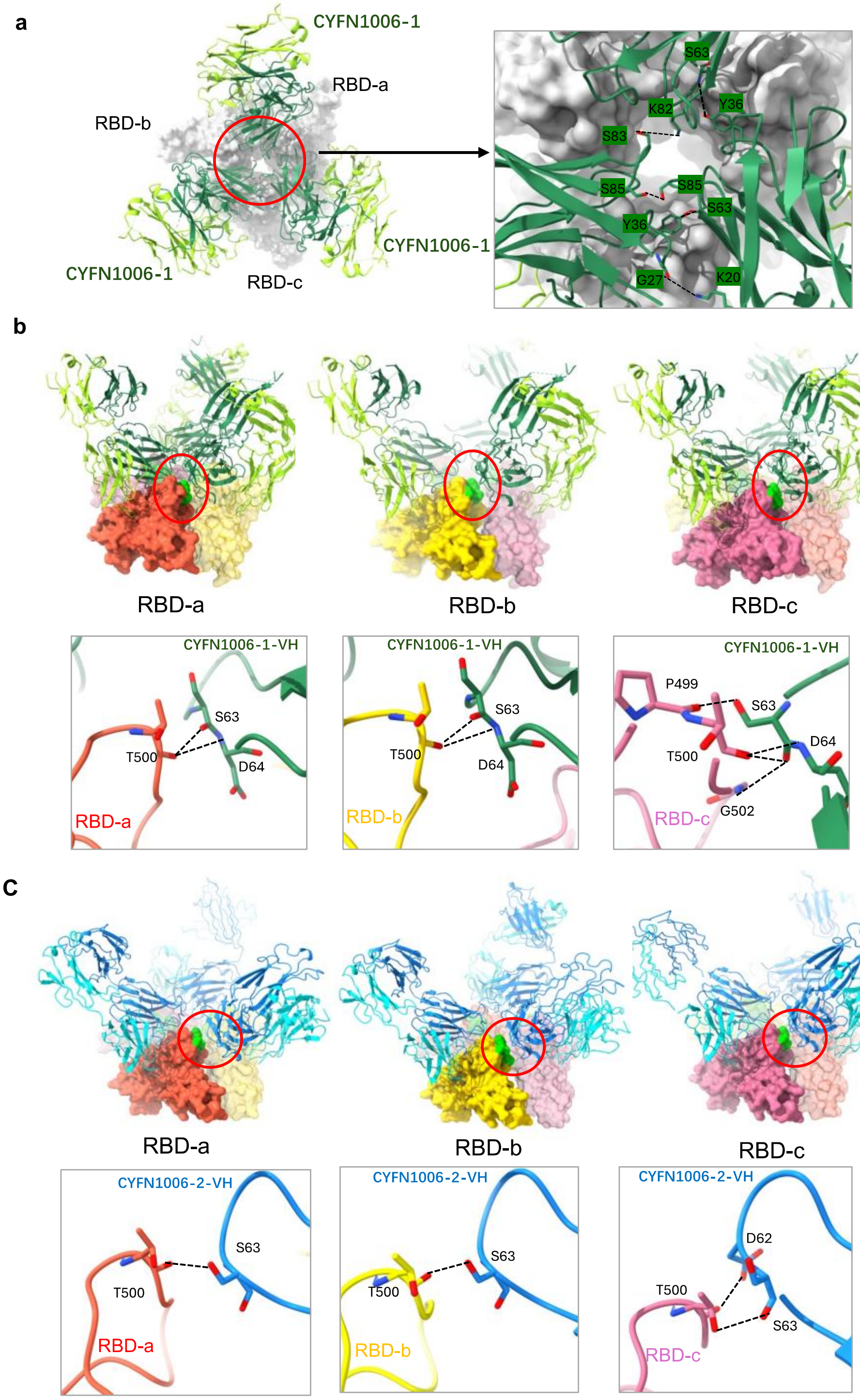


**Extended Data Fig. 7 Details of the interaction of CYFN1006-1/CYFN1006-2 with adjacent fabs and RBDs.**

(**a**) A top view of the complex of the XBB.1.16 RBDs with CYFN1006-1 Fabs and the interactions between the three adjacent heavy chains. Hydrogen bonds are represented by dashed lines. (**b**) Interactions between XBB.1.16 RBD and the neighboring

CYFN1006-1 heavy chain. Three RBDs are shown as orange, yellow, and pink surfaces respectively, and the RBD epitopes involved in the interaction are colored green. Hydrogen bonds are represented by dashed lines. (**c**) Interactions between EG.5.1 RBD with neighboring CYFN1006-2 heavy chain. Hydrogen bonds are represented by dashed lines.

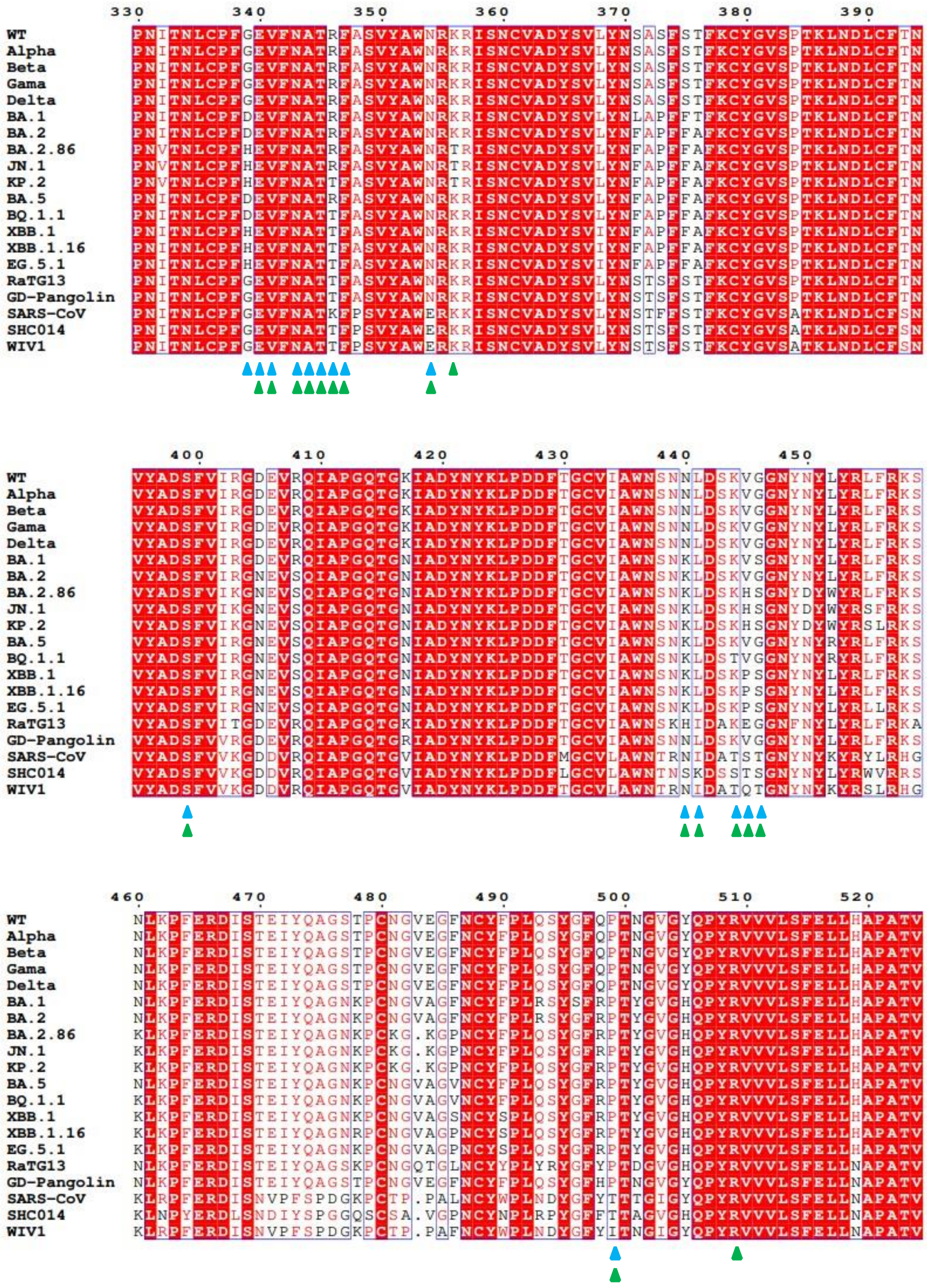

**Extended Data Fig. 8 Sequence alignment of SARS-CoV-2 variants.**

Conserved amino acids are highlighted as red. Residues involved in the binding of CYFN1006-1 and CYFN1006-2 are labeled green and blue, respectively.

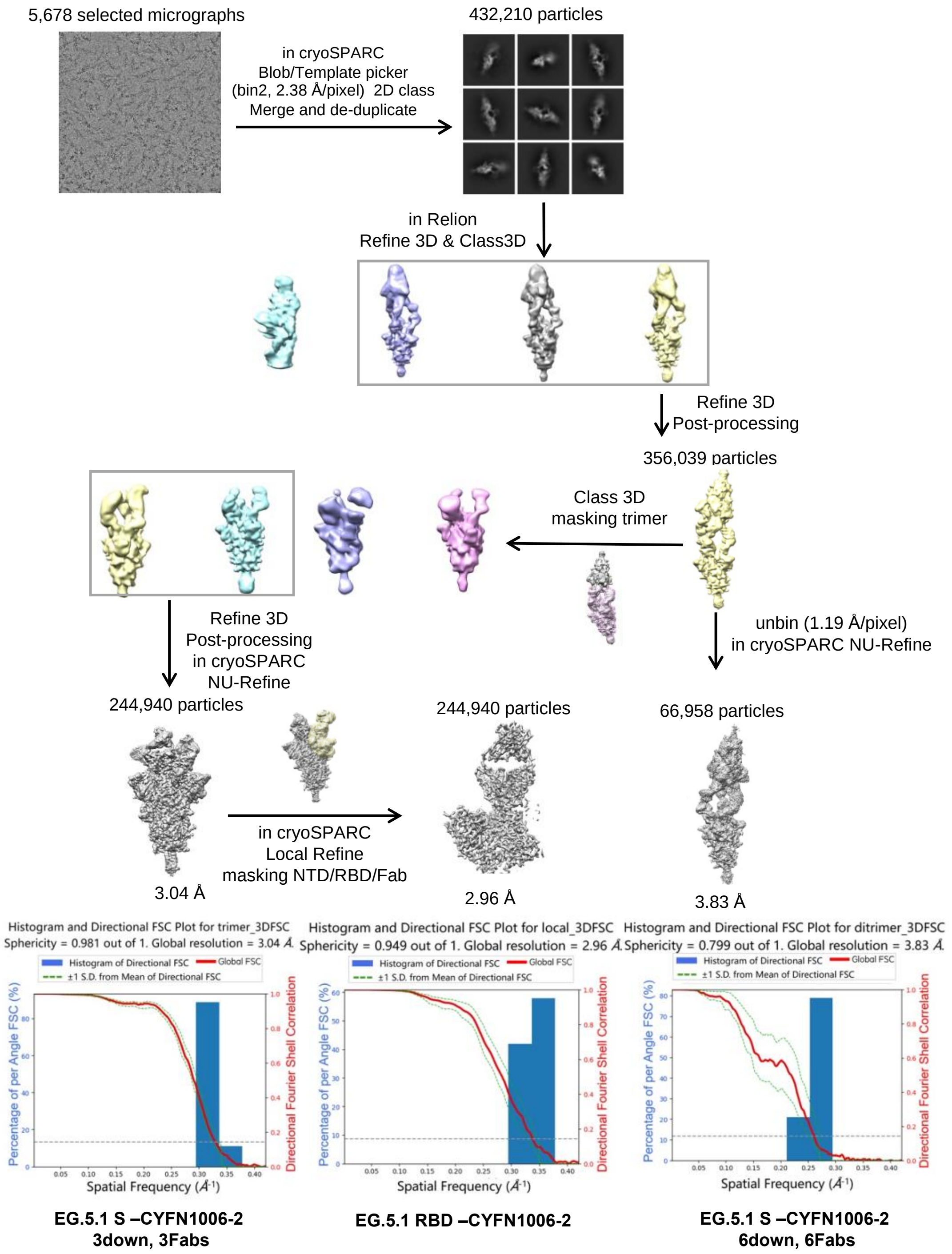


**Extended Data Fig. 9 Data processing flowchart of EG.5.1 S-CYFN1006-2 complex.**

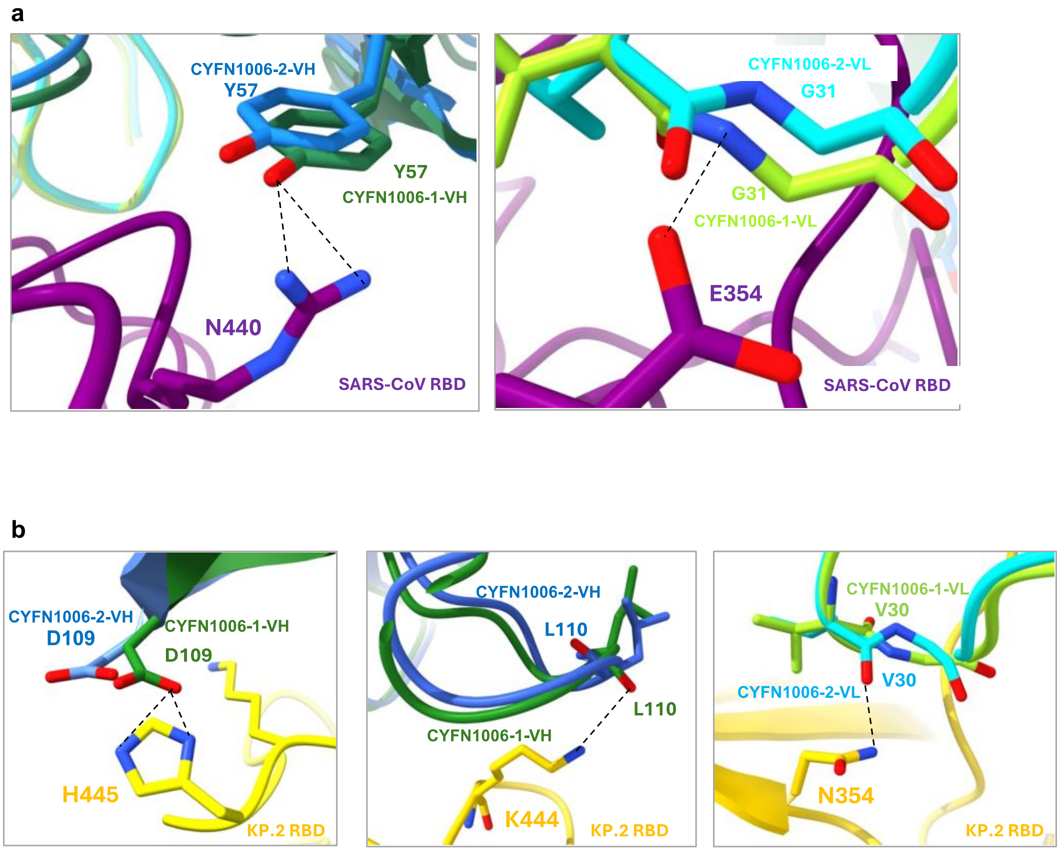


**Extended Data Fig. 10 Prediction of interactions between CYFN1006-1/CYFN1006-2 and SARS-CoV S/KP.2 S.**

**(a)** SARS-CoV RBD (PDB:2DD8) is represented as a purple ribbon, the different hydrogen bonds of the two antibodies are represented as dashed lines. Hydrogen bonds are represented as dashed lines. **(b)** KP.2 RBD is represented as a yellow ribbon, the different hydrogen bonds of the two antibodies are represented as dashed lines. Hydrogen bonds are represented as dashed lines. KP.2 RBD model is generated using Swiss-model.

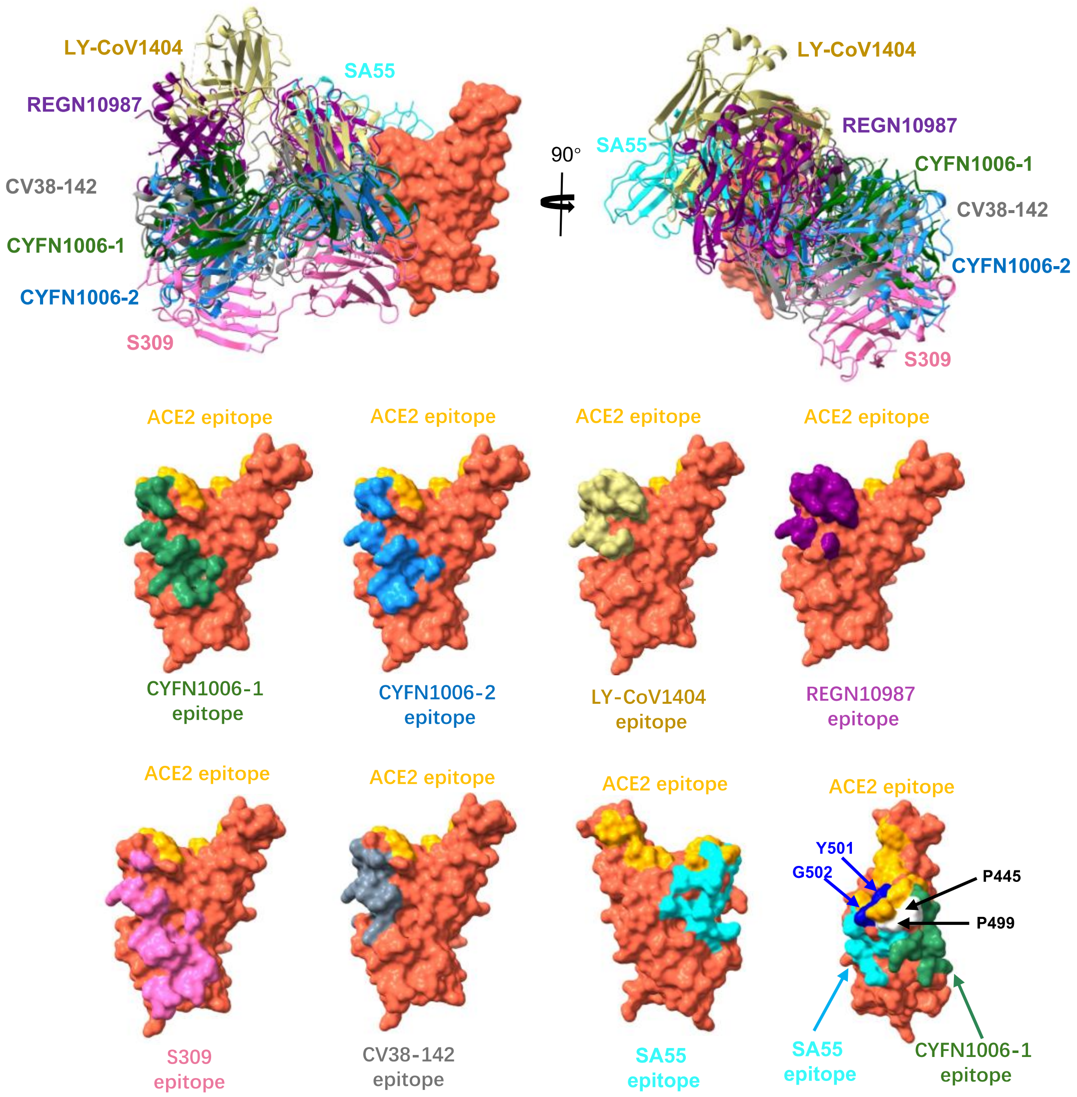


**Extended Data Fig. 11 Epitope comparison of similar antibodies on the RBD of SARS-CoV-2 S.**

(**a**) Comparison of the CYFN1006-1 (green), CYFN1006-2 (dodger blue), LY-CoV1404 (PDB:7MMO, pale goldenrod), REGN10987 (PDB:6XDG, purple), S309 (PDB:7XCK, pink), CV38-142(PDB:7LM8, gray) and SA55 (PDB:7Y0W, red) epitopes on RBD. (**b**) Surface representation of RBD showing the buried regions by CYFN1006-1, CYFN1006-2, LY-CoV1404, REGN10987, S309, CV38-142 and SA55, respectively. The blue region represent the overlap of SA55 and ACE2 epitopes, and the white region represents the overlap of SA55 and CYFN1006-1 epitopes.

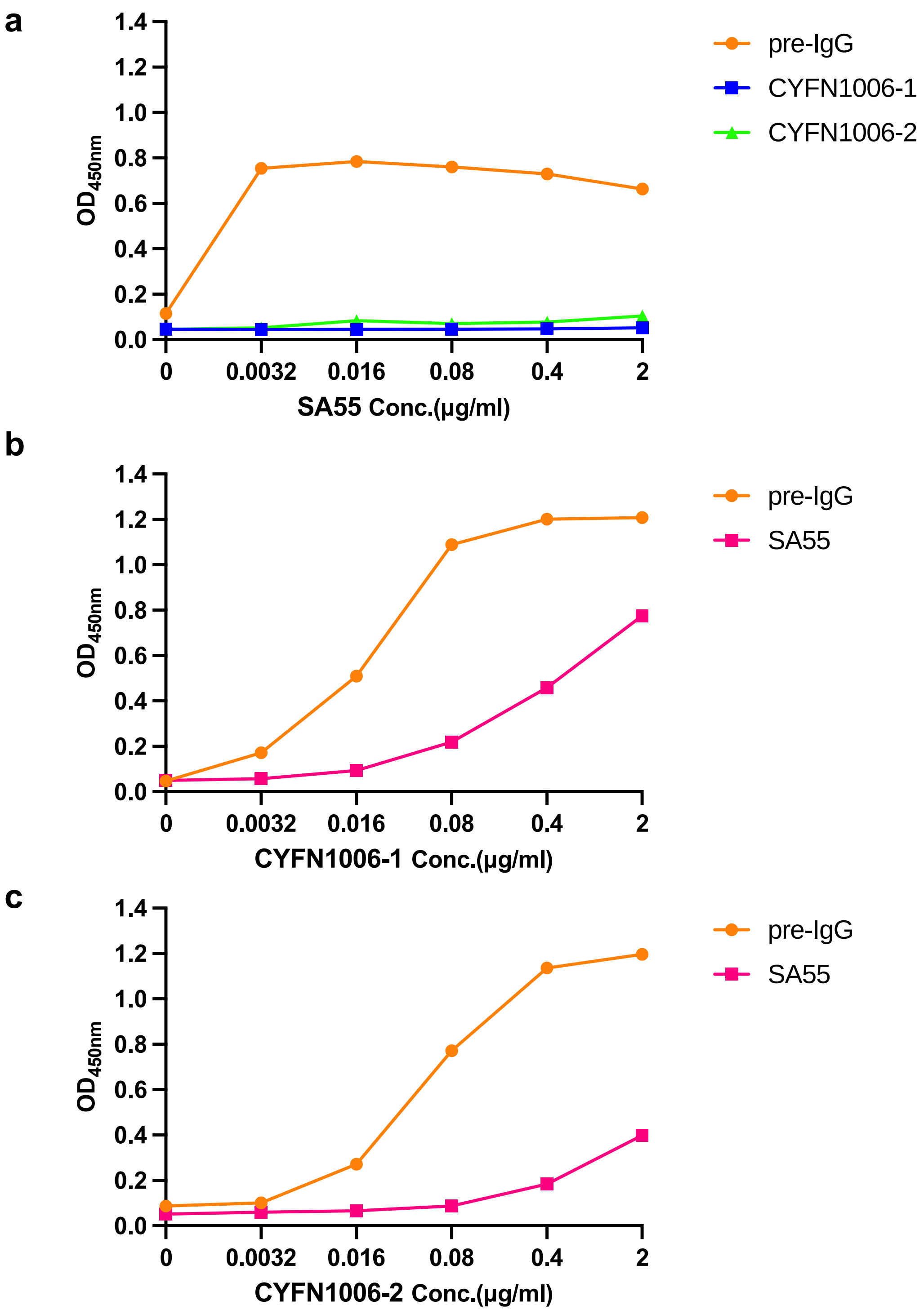


**Extended Data Fig. 12 Competitive binding curves of CYFN1006-1, CYFN1006-2 and SA55 to SARS-CoV-2 RBD.**

(**a**) CYFN1006-1 or CYFN1006-2 fully competes SA55 for binding to RBD. (**b,c**) SA55 partially competes CYFN1006-1 or CYFN1006-2 for binding to RBD.
